# Bacterial diets differentially alter lifespan and healthspan trajectories in *C. elegans*

**DOI:** 10.1101/2020.04.29.068288

**Authors:** Nicole L. Stuhr, Sean P. Curran

## Abstract

Diet is one of the more variable aspects in life due to the variety of options that organisms are exposed to in their natural habitats. In the laboratory, *C. elegans* are raised on bacterial monocultures, traditionally the *E. coli B* strain OP50, and spontaneously occurring microbial contaminants are removed to limit experimental variability because diet - including the presence of contaminants, can exert a potent influence over animal physiology. In order to diversify the menu available to culture *C. elegans* in the lab, we have isolated and cultured three such microbes: *Methylobacterium, Xanthomonas*, and *Sphingomonas.* The nutritional composition of these bacterial foods is unique, and when fed to *C. elegans*, can differentially alter multiple life history traits including development, reproduction, and metabolism. In light of the influence each food source has on specific physiological attributes, we comprehensively assessed the impact of these bacteria on animal health and devised a blueprint for utilizing different food combinations over the lifespan, in order to promote longevity. The expansion of the bacterial food options to use in the laboratory will provide a critical tool to better understand the complexities of bacterial diets and subsequent changes in physiology and gene expression.

## INTRODUCTION

Since the discovery that aging can be improved by dietary, genetic, and pharmacological interventions, there has been an increase in studies aiming to elucidate how each of these interventions can promote long life and healthy aging. Many dietary interventions including calorie restriction, intermittent fasting, and dietary restriction^1^ have been shown to not only increase maximal lifespan, but average lifespan of the cohort and healthspan as well. New diets and fads have become popularized within society^2,3^, however, these dietary fads focus on removing one aspect of diet (carbohydrates, sugars, fats, proteins, etc.) instead of focusing on the overarching complexity of food.

Food is imperative for all organisms in order to provide nourishment to fuel growth and essential cellular functions. Diet is one of the more variable aspects in life due to the vast options that organisms are regularly exposed to in their natural habitats. Survival within these environments require organisms to select high quality food sources from a range of nutritiously diverse alternatives. For centuries, society has accepted that diet is important for health and longevity; “You are what you eat.” However, our understanding of why diet holds such profound influence over our health and how we can use this knowledge to improve overall health and longevity requires further study.

Many diet studies have employed the use of the model organism *Caenorhabditis elegans*^4–6^ due in part to the many shared core metabolic pathways with mammals^6^. Understanding how the introduction of specific dietary foods impact both health and lifespan in the worm may aid in explaining the variability in the rates of aging and severity of age-related disease. These bacterivore nematodes can be used for the identification of dietary effects due to their invariant and short developmental and reproductive periods, which are followed by an averaged 3-week lifespan^7^.

The impact of bacterial diets on physiology is vast and certainly integrates into multiple life history traits including development^8^, reproduction^9^, healthspan^10^, and longevity^4,11–13^. Additional studies have identified diet-induced phenotypic effects that are accompanied by measurable differences in metabolic profiles^11,14,15^, fat content ^5,6,8,16^, and feeding behavior^8,13,17,18^. *C. elegans* interact with a variety of bacterial species in their natural environment, including Proteobacteria, Bacteroidetes, Firmicutes, and Actinobacteria^19^. Under normal laboratory conditions, *C. elegans* are cultured using the standardized bacterial species *Escherichia coli* OP50. This bacterium was not chosen because of its association with nematodes in the wild, but because of availability in the laboratory setting^16^. Maintenance of *C. elegans* in the laboratory includes the removal of spontaneously occurring bacterial contaminants in order to limit experimental variations. The lack of exposure of *C. elegans* to naturally occurring bacterial species have led to many aspects of *C. elegans* biology becoming undetectable in the artificial laboratory environment^16,19–22^.

We routinely noticed that *C. elegans* are found feeding on contaminants, when present, rather than the *E. coli* food source supplied. With this in mind, we looked at these contaminants as an opportunity to examine the impact that different food sources have on physiology. We identified the genera of three such “contaminants” as bacterium that can be found in *C. elegans* natural habitats. As such, we decided to use these three microbes alongside the three most commonly used *E. coli* fed to worms, to expand the menu of bacterial diets and provide insight into the contribution that different foods have on *C. elegans* physiology^4,23^. Our studies provide a comprehensive assessment of changes in physiology and transcriptomic signatures as a result of *C. elegans* exposure to bacteria found in the laboratory environment versus their natural environment. To our knowledge, this is the first side-by-side comparison documenting how these six bacteria differentially effect multiple key aspects of *C. elegans* physiology over the lifespan. These data represent a critical resource to realize the impact different bacterial foods can have on multiple aspects of *C. elegans* physiology and gene expression.

## RESULTS

### Identification and characterization of three distinct bacterial diets

In their natural environment, *C. elegans* need to adapt to a variety of microbes they might encounter as a source of food. The necessity to cope with varied food sources have likely shaped multiple aspects of their biology, which are masked in the artificial laboratory environment where they are fed the unnatural diet of *E. coli*^7^. In order to elucidate the underlying connection between bacterial diet and physiology, it is important to evaluate how different foods influence behavior and physiological attributes.

We isolated and cultured three bacterial species as noteworthy *C. elegans* foods that we call “Red”, “Orange”, and “Yellow” due to their pigments when grown on plates (**Fig. 1a**). Using 16S ribosomal sequence alignment, we identified the genus of each new bacterial diet: Red is *Methylobacterium*; Orange is *Xanthomonas*, and Yellow is *Sphingomonas.* Intriguingly, these three bacterial diet genera can be found in *C. elegans* natural environments^16^. We next identified optimal growth parameters to ensure monoculture growth for each bacteria in LB broth and on plates (Supplementary Fig. 1a). Although most species were able to grow at 37°C on LB solid media and in LB liquid media, Red/*Methylobacterium* required a growth temperature of 30°C on LB solid media, while Yellow*/Sphingomonas* required a growth temperature of 26°C in LB liquid media. The growth rate of each bacterial species was similar, except Red/*Methylobacterium*, which was markedly reduced (**Fig. 1b** and Supplementary Fig. 2a).

**Figure 1.**
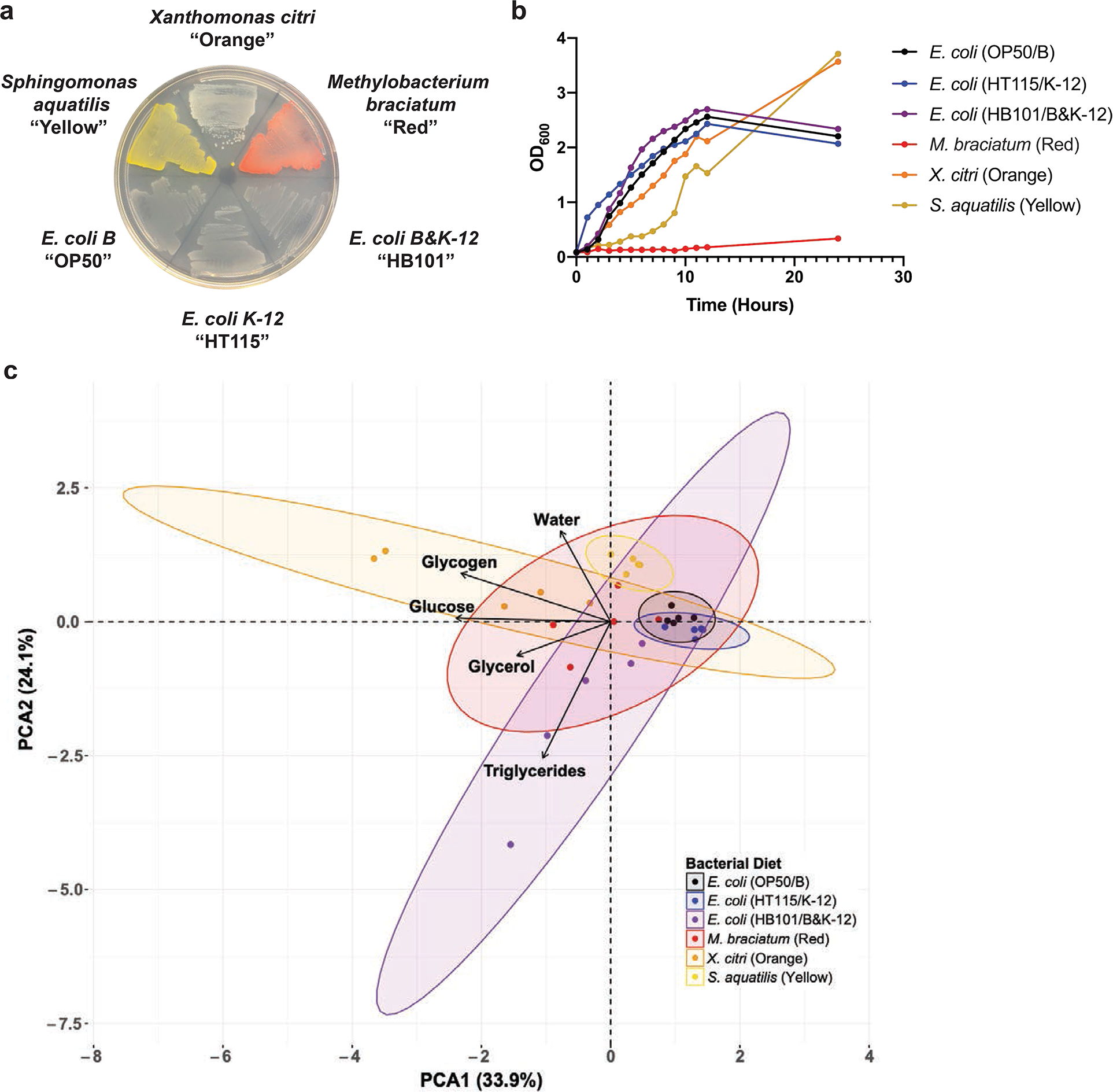
Characterization of bacterial diets. (a) Bacterial streak plate with the six bacterial diets fed to *C. elegans*. (b) Growth curves of the bacteria with antibiotics, with optical density measurements every hour for 12 hours and another measurement taken at 24 hours. (c) Principal component analysis (PCA) of metabolite concentrations in the different bacterial diets. Bacteria were collected during the log phase of growth for bomb calorimetry (water content) and metabolite kits.

Preceding research has identified many distinct differences in macronutrient composition between the three *E. coli* strains used to raise *C. elegans* in the laboratory. Not only do these strains differ in carbohydrate levels 24,25, but also contain varying levels of vitamin B12 ^25^, dietary folate and tryptophan ^26–28^. The dissimilarity between these bacterial compositions has been shown to alter many phenotypic attributes in *C. elegans*. To assess the nutritional value of the bacterial diets used in our studies, we first performed bomb calorimetry to define the total caloric composition of each bacteria. Surprisingly, the total caloric composition was not significantly different in any of the bacteria in comparison to the *E. coli* OP50/B bacterial diet. In order to better assess the differential nutrient composition of each bacteria, we next measured the concentrations of glucose, glycerol, glycogen, triglyceride, and water to define a nutritional profile for each bacterial food source (**Fig. 1c** and Supplementary Fig. 1b-g). Orange/*Xanthomonas* was the most significantly different food among the three distinct bacterial diets because glucose, glycerol, glycogen, and water content were all higher than the levels found in OP50/B. Red/*Methylobacterium* displayed a similar trend with an increase in glycerol, glucose, and water content, but not triglycerides. Finally, Yellow/*Sphingomonas* only carried more glycerol and water content, compared to *E. coli* OP50/B. Interestingly, *E. coli* HB101/B&K-12 was the only bacterial diet with an increase in triglyceride content relative to *E. coli* OP50/B. We also noted that glucose and glycogen content were highest in Orange/*Xanthomonas*, glycerol and triglyceride content were highest in *E. coli* HB101/B&K-12, and water content was highest in Yellow*/Sphingomonas.* Taken together, Red/*Methylobacterium*, Orange/*Xanthomonas*, and Yellow/*Sphingomonas* represent three new potential *C. elegans* food sources, with similar calorie content, but differential nutritional composition to *E. coli* OP50/B that could be used to investigate dietary effects on animal physiology.

### Bacterial diets direct unique transcriptional signatures

We next confirmed that worms could be cultured on each bacteria as the sole source of nutrition to enable the systematic characterization of the impact of these foods on *C. elegans* health. After maintaining animals on each bacterial food for more than 30 generations, we then compared animals reared on these new laboratory bacterial foods alongside populations grown on the three most common *E. coli* foods (OP50, HT115 and HB101).

Because diet can impact multiple cellular processes, we first assessed the steady state transcriptional response to each bacterial diet by RNA-seq (**Fig. 2a** and Supplementary Data 1). Each food source evoked a unique transcriptional signature (**Figs. 2b-f** and Supplementary Data 1) when fed to the animal over multiple generations, which revealed the impacts of food on multiple physiological responses. We compared the relative expression for each gene to age-matched animals fed OP50 and discovered that there were 743 genes altered in animals fed HT115, 3512 genes altered in animals fed HB101, 455 genes altered in animals fed Red, 1149 genes altered in animals fed Orange, and 6033 genes altered in animals fed Yellow. Of these, 102 genes were uniquely altered in animals fed HT115, 320 on HB101, 40 on Red, 24 on Orange, and strikingly 2667 on Yellow (**Fig. 2g** and Supplementary Data 1). In addition to finding genes that were shared on two, three, and four food types, we also identified 140 genes that were altered on all five bacterial foods, as compared to OP50-fed animals. With the analysis of these 140 genes, we were able to see that in terms of transcriptional response, Red and HT115 were more similar to each other and more closely related to OP50 while Orange, Yellow, and HB101 shared many similarities (**Fig. 2h**).

**Figure 2.**
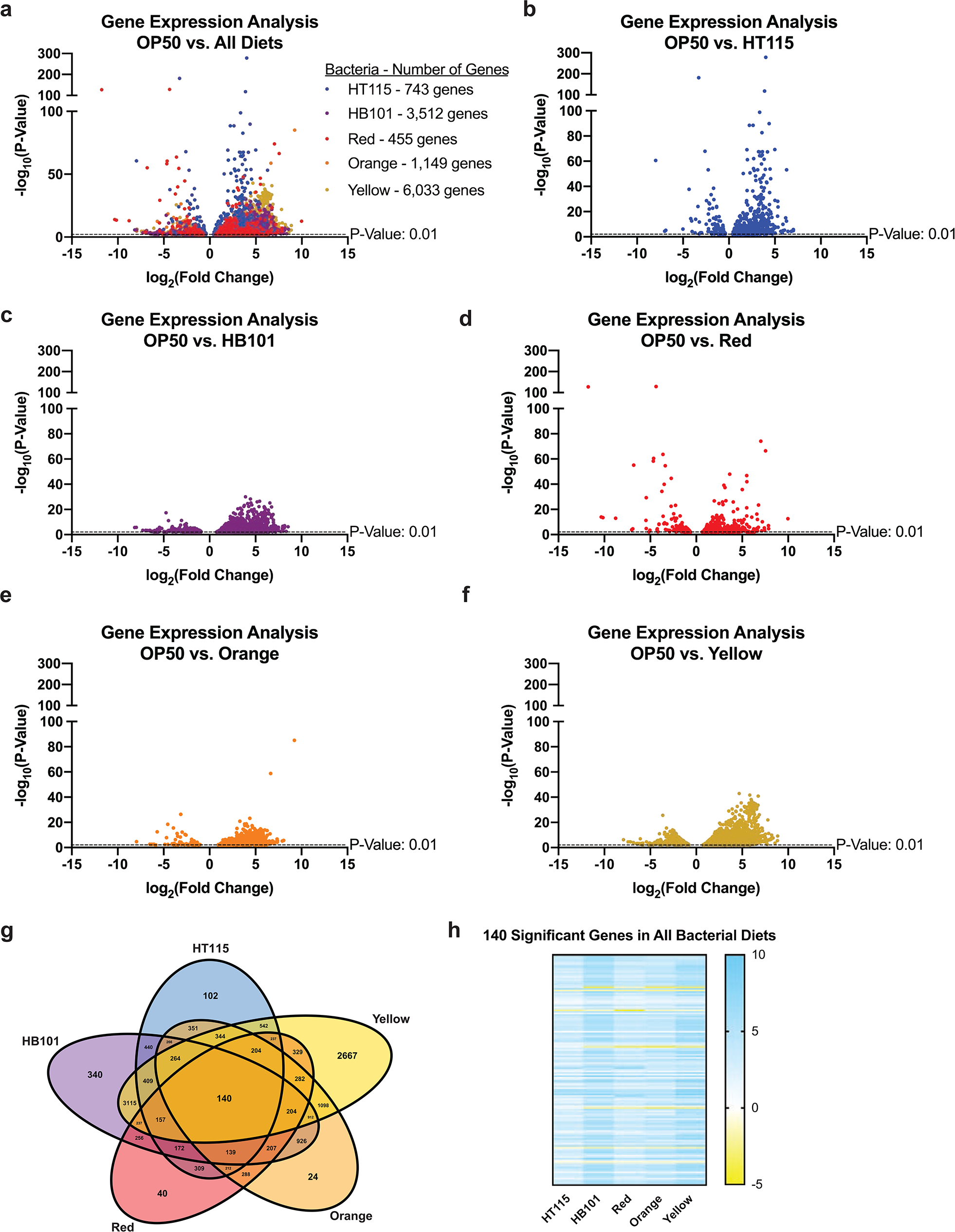
Gene expression analysis of L4 *C. elegans* on each bacterial food source after thirty generations. (a) Volcano plot of all differentially expressed genes in each food relative to OP50-reared worms. All genes considered to be significant have a p-value <0.01. (b-f) Volcano plots of each individual food with all significant genes that are differentially expressed. (g) Venn diagram of all significant genes. Shows the number of genes shared between two, three, four and five of the bacterial diets, along with the number of genes unique to each bacterium. (h) Heat map contains genes that are shared between all five bacterial diets. Gene Ontology (GO) terms for uniquely-expressed genes are noted below and split between genes that were up-regulated and down-regulated. More information for the GO-term enrichment analysis of RNAseq data in larval stage 4 animals can be found in Supplementary Data 1 and 2.

An analysis of the Gene Ontology (GO)-terms of the unique genes that were differentially expressed in animals fed each bacteria type revealed specific molecular signatures for each diet (Supplementary Table 1). Notably, we observed expression changes for genes that could alter multiple physiological processes including development, metabolism, reproduction, and aging. In consideration of this enrichment, we explored the impact of the different bacterial food sources on each of these critical physiological attributes.

### Food-induced acceleration of developmental timing

Our ability to culture animals on each bacterial food revealed that they were sufficient to support life, but as food can influence multiple life history traits, we further examined the physiological responses to the foods relative to the standard OP50 food. Bacterial diet can impact developmental timing^24^ and as such, we asked whether any bacterial food altered the time to reach each larval stage using the *mlt-10*::GFP-PEST reporter strain, which marks the transition of each animal as it progresses through its four developmental molts^29^; noting that on OP50 larval development occurs with invariant timing (**Fig. 3a**). We determined that all bacterial foods facilitated successful development to reproductive adulthood but remarkably, each food source resulted in faster development relative to OP50-reared animals (**Figs. 3b-f** and Supplementary Fig. 3). We noted three important variables in this precocious development: first, bacteria could accelerate development of specific larval stages – HT115 (**Fig. 3b**) and Red (**Fig. 3c**) resulted in early transition from larval stage 2 (L2) to larval stage 3 (L3), Orange (**Fig. 3d**) resulted in early transition from L3 to larval stage 4 (L4), and HB101 (**Fig. 3e**) and Yellow (**Fig. 3f**) resulted in early transition from L4 to adulthood. Second, with the exception of the precocious transition between two larval stages listed above, the time spent in each larval stage (peak to peak time) was one to three hours shorter compared to OP50 raised animals (Supplementary Fig. 3). Third, the time to molt (trough to trough time) was mostly unchanged (**Fig. 3a**). In addition, we measured animal size at each developmental stage, which showed some small but significant differences in body size on different bacterial foods, similar to observations from other studies^30–33^. Worms raised on HT115 contained a bigger area from L4-day 1 adult stage, Orange worms had a smaller area until day 2 of adulthood, and the other foods resulted in a fluctuation of larger and smaller areas throughout developmental stages, relative to OP50 (Supplementary Fig. 3). These data reveal that animals fed the new Red, Orange, and Yellow bacterial diets or the commonly used *E. coli* foods HT115 or HB101, reach reproductive maturity faster, as compared to the standard OP50 food source, which could influence animal fitness.

**Figure 3.**
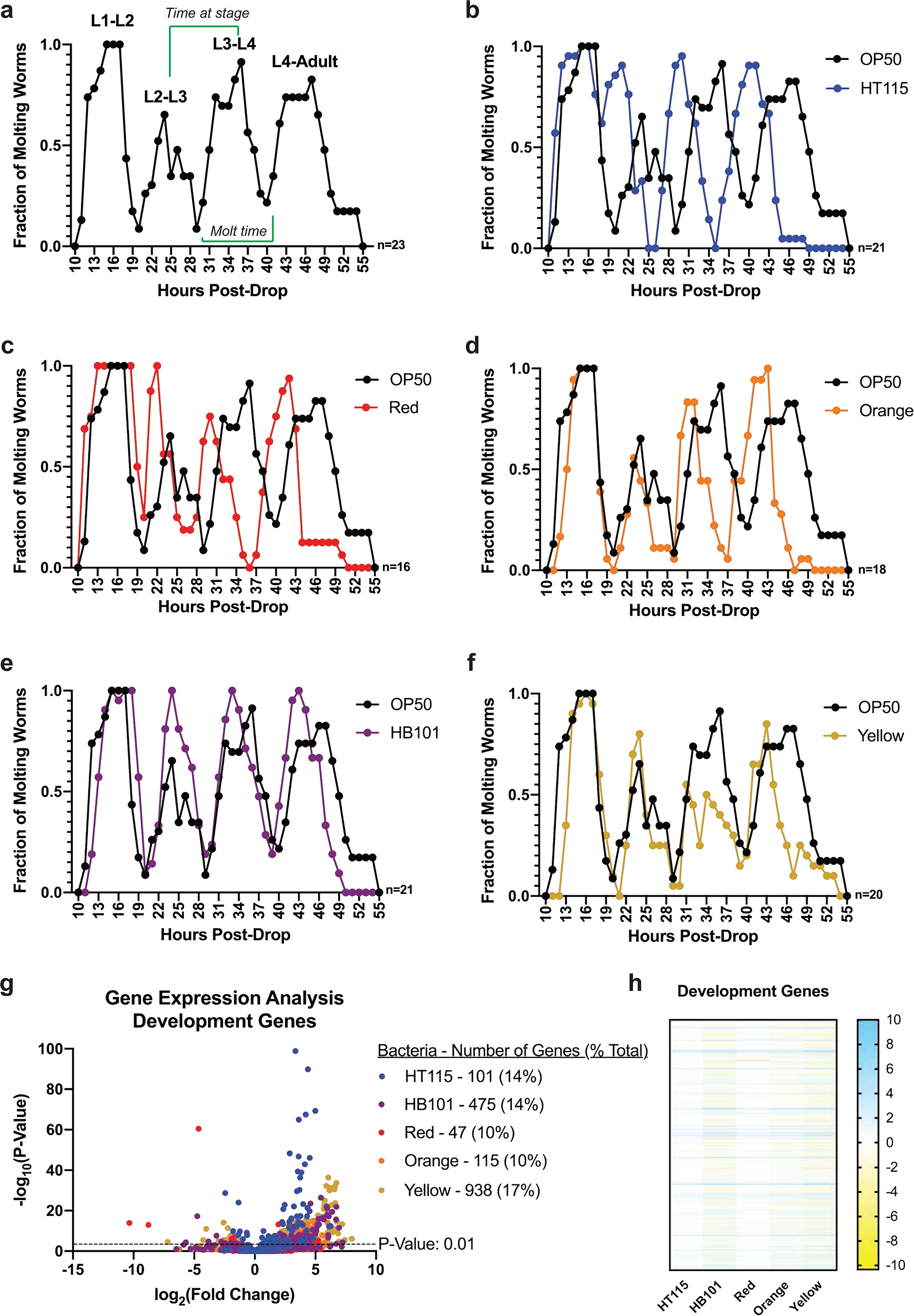
Developmental timing of *C. elegans* is dependent upon bacterial diet. (a) The molting reporter strain *mgls49[mlt-10p::gfp-pest ttx-3p::gfp] IV* allows for the visualization of molting between larval stages of development into adulthood. The time between peaks represents the period at each larval stage, while time between troughs represent time spent molting. Analysis was carried out after this strain was raised on each bacterial food for twenty generations. (b-f) Developmental timing of worms raised on each food relative to OP50. All diets showed accelerated development into adulthood when compared to worms raised on OP50. (g) Volcano plot showing how many of the differentially expressed genes are related to development. The number of genes on each bacterial food are labeled in the legend. All genes have a p-value <0.01. (h) Heat map of development genes that are significant with a p-value <0.01 in at least one of the five bacterial diets, relative to the OP50 bacterial diet.

Due to both the difference in total time to reproduction and time to progress through each developmental stage, we examined the expression of *C. elegans* genes important for development (**Figs. 3g-h)**. Based on GO-term classifications, worms fed the Yellow bacteria had the greatest number of differentially expressed developmental genes, while worms fed Red bacteria had the fewest. Intriguingly, of the 140 genes with shared expression changes between the five diets, 12 of these genes are involved in the molting cycle (*sqt-2,sqt-3,ptr-4,ptr-18,mlt-7,mlt-10,qua-1,bli-1,dpy-3,rol-6,acn-1* and *noah-1*)^29,34^.

### Bacterial diets differentially affect lipid homeostasis

Diet directly impacts overall metabolism, which is under tight genetic control^23,35^. The unique metabolic profiles of each bacteria drove us to explore how these different microbial diets could affect organismal metabolism. As changes in intracellular lipid stores are easily measured by microscopy, we stained age-matched populations of fixed animals with two lipophilic dyes: Nile Red (NR) to measure total lipid content (**Figs. 4a-g**) and Oil Red O (ORO) to identify lipid distribution (**Figs. 4h-n** and Supplementary Fig. 4). Relative to OP50-fed animals (**Fig. 4a**), overall lipid levels were increased in animals fed HT115 (**Figs. 4b,g**) and Orange (**Figs. 4e,g**), but were markedly decreased in HB101 (**Figs. 4c,g**), Red (**Figs. 4d,g**), and Yellow (**Figs. 4f-g**). Under most bacterial foods, lipid distribution across tissues were similar (**Figs. 4h-l,n** and Supplementary Fig. 4), except for animals fed the Yellow bacteria, which resulted in the depletion of intestinal lipids in late reproductive adults, a phenotype known as age-dependent somatic depletion of fat (Asdf) at day 3 of adulthood (**Figs. 4m-n**)^36,37^.

**Figure 4.**
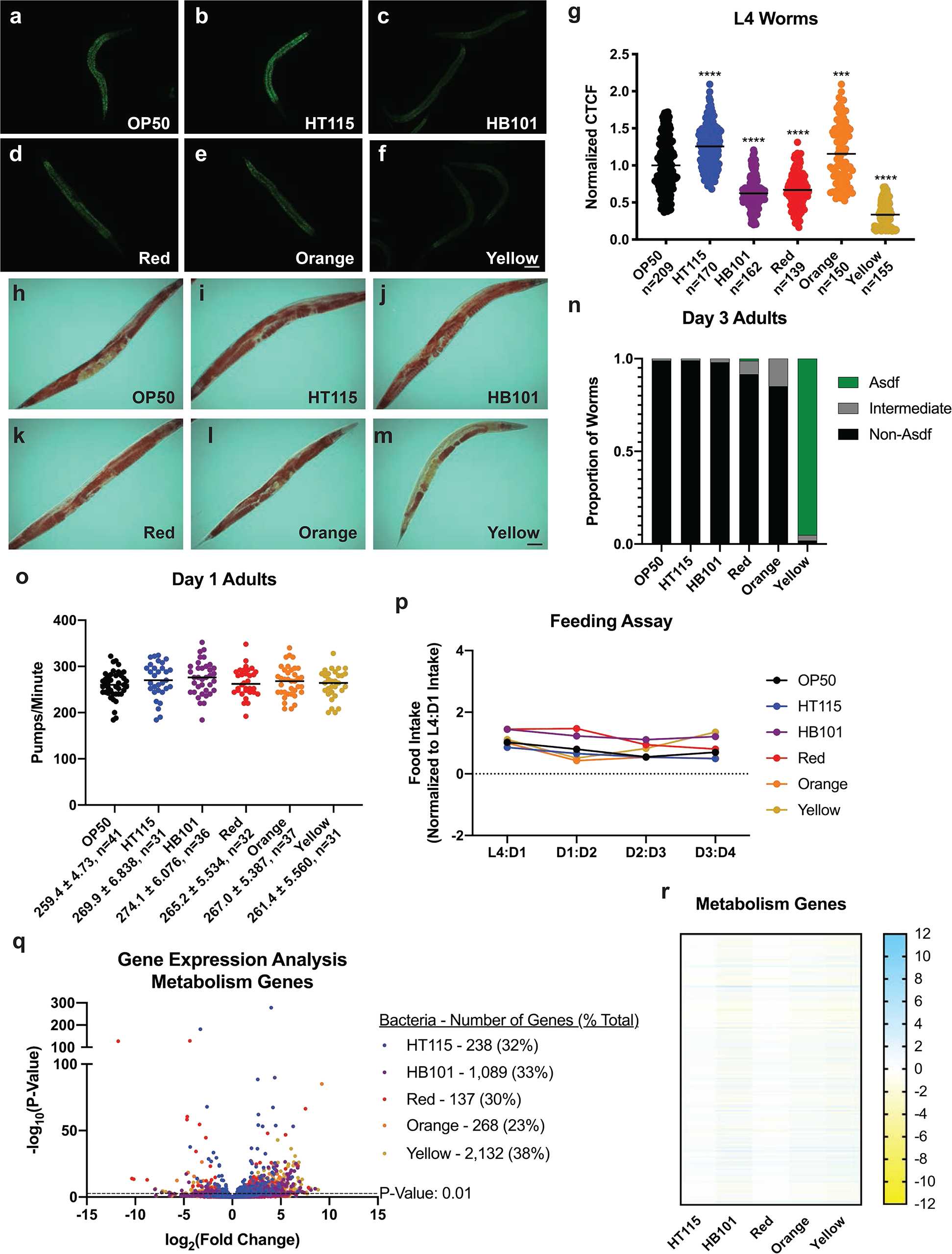
Fat content and distribution vary depending on the food *C. elegans* are exposed to. (a-f) Nile Red staining of L4 worms, scale bar 50 μm. HT115 (b) and Orange (e) have significantly higher fat content while HB101 (c), Red (d) and Yellow (f) have lower fat contents. (g) Quantification of Nile Red staining with comparisons made to the OP50 control. GFP fluorescence is measured and then normalized to area before being normalized to OP50. Statistical comparisons by Tukey’s multiple comparison test. ***, p<0.001; ****, p<0.0001. (h-m) Oil Red O lipid staining in day 3 adult *C. elegans*. OP50, HT115, HB101, Red, and Orange show similar lipid distribution of fat throughout the animal (representative image, scale bar 50 μm). (m) Yellow-reared worms display age-dependent somatic depletion of fat (Asdf) with loss of fat in the intestine while fat is retained in the germline (representative image). (n) Quantification of lipid distribution across tissues. (o) Pumping in day 1 adult worms is not significantly different between any of the bacterial foods and OP50. (p) Food intake of *C. elegans* raised on each bacterial food in liquid nematode growth media. (q) Volcano plot of differentially expressed metabolism genes in all bacterial diets compared to OP50. All genes have a p-value <0.01. (r) Heat map of metabolism genes that are significant with a p-value <0.01 in at least one of the five bacterial diets, relative to the OP50 bacterial diet. All studies were performed in biological triplicate.

The fact that animals fed each of the bacteria developed faster than OP50-reared animals indicates animals were not nutritionally deprived, which can result in developmental delay^19,38–42^. Nevertheless, different microbes have been shown to alter rates of pharyngeal pumping, which regulates ingestion of bacteria^43^. Pharyngeal pumping rates were examined in both the L4 stage and day 1 adult animals, but none of the bacterial diets significantly altered pharyngeal pumping (**Fig. 4o** and Supplementary Fig. 4). As a complement to the pharyngeal pumping analyses, we wanted to measure overall food intake from the L4 stage of development to day 4 of adulthood (**Fig. 4p**). Similar to the pharyngeal pumping data, the food intake assay also demonstrated that the amount of food being eaten by a single worm at a certain point in development was not significantly different. This indicates that worms are eating similar amounts of food throughout their lives, which suggests that fat phenotypes observed above are not due to the amount of food ingested, but rather the composition and nutritional aspect of the bacteria.

Because each bacterial diet represented a specific metabolic profile and *C. elegans* reared on these bacteria differed in the amounts of stored intracellular lipids, we examined the expression of genes that regulate lipid metabolism and homeostasis (**Figs. 4q-r**). Of the total number of genes that were differentially expressed on each bacterial food source, about 1/3 of the genes in worms raised on each food were related to metabolism. Feeding animals the Yellow bacteria evoked the largest number of metabolism-related gene changes, which correlated with our phenotypic observation that animals fed this food store the lowest levels of fat at the L4 stage (**Fig. 4g**) and it was the only bacteria to drive the Asdf phenotype later in life (**Fig. 4m**). Among the genes differentially expressed on these new food sources were *fat-5* and *fat-7*, which may suggest changes in lipid biosynthesis pathways, as well as multiple lipases (Lipl-class), suggesting changes in lipid utilization (**Fig. 4g**).

### Red and Yellow reduce reproductive output

We next investigated how each bacterial diet influenced fertility by measuring the total number of viable progeny laid by individual animals over a ten-day reproductive span. The total number of progeny was reduced by ~25% when animals were fed the Red bacteria, modestly reduced (~10%) in animals fed the Yellow bacteria, and unchanged on the other microbial foods (**Fig. 5a**). Although animals reached peak reproductive output at approximately the same time, animals fed Red and HB101 ceased reproduction sooner than all other groups (**Fig. 5b** and Supplementary Fig. 5). Reproductive output of *C. elegans* hermaphrodites is determined by the number of spermatids generated at the L4 larval stage^44,45^. To determine if the decreased progeny production and early loss of reproductive output were due to diminished sperm availability in Red-reared worms, we mated hermaphrodites with males, which increases total reproductive output (**Fig. 5c**). Males raised on either OP50 or Red could increase total reproductive output similarly when mated to OP50-reared or Red-reared hermaphrodites, which indicated that sperm are functional when in excess. Surprisingly, when mated, hermaphrodites raised on Red have significantly more progeny as compared to hermaphrodites raised on OP50. We also counted the number of unfertilized oocytes laid by unmated hermaphrodites, which revealed HT115, HB101, Red, and Orange bacteria resulted in markedly reduced expulsion of unfertilized gametes (**Fig. 5d**) while hermaphrodites on Yellow were similar to animals fed OP50. An analysis of the reproduction-related genes that were differentially changed in animals fed each of the bacterial foods revealed several bacteria-specific changes (**Figs. 5e-f**).

**Figure 5.**
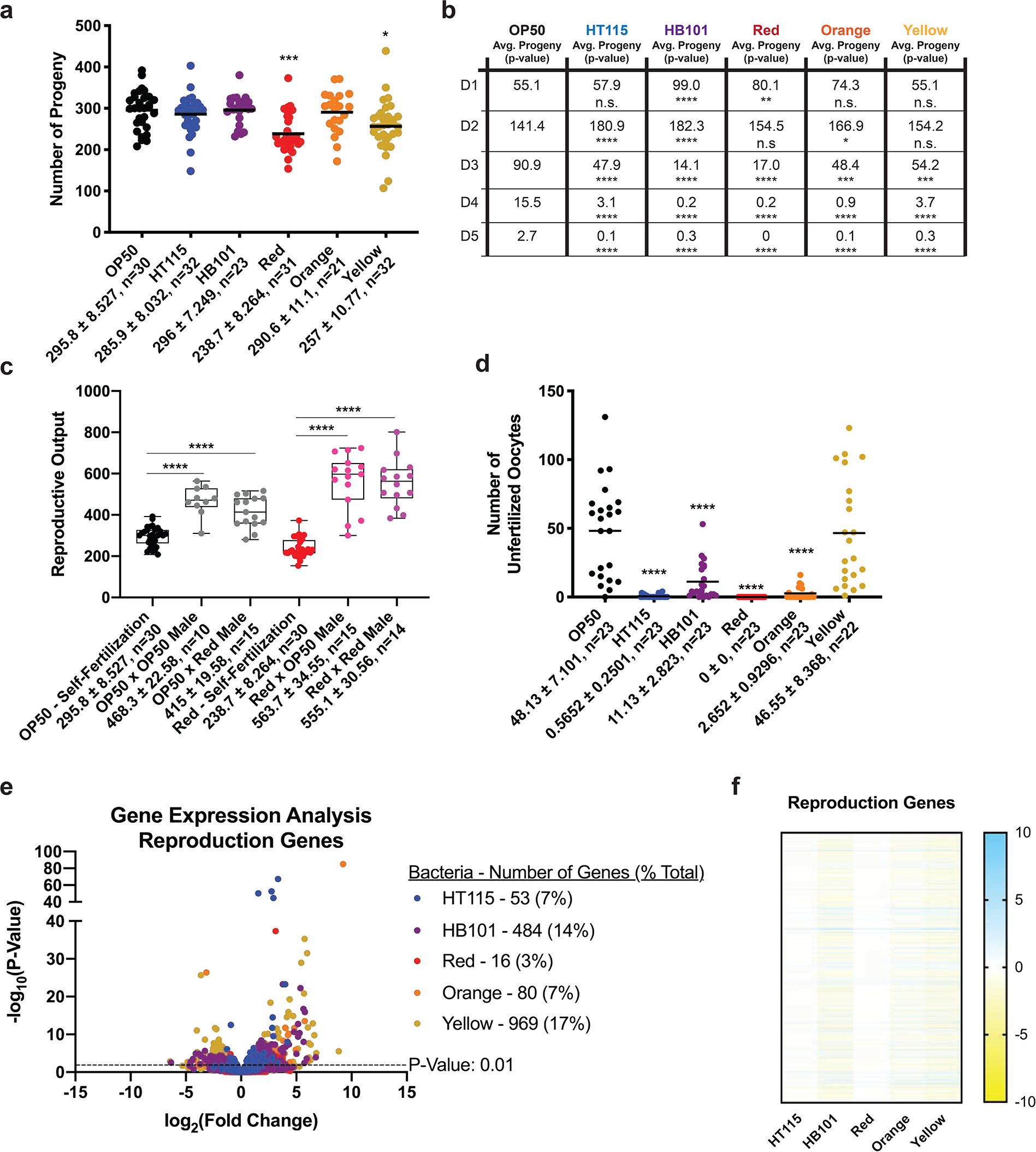
*C. elegans* raised on Red and Yellow have decreased reproductive output. (a) Hermaphrodites have reduced brood size when fed Red or Yellow bacteria. (b) Reproductive timing is also altered, revealing differences between all bacterial diets and OP50 in total output each day of their reproductive span. HB101 and Red halt output of viable progeny before OP50, HT115, Orange and Yellow. (c) OP50 and Red hermaphrodites mated with OP50 or Red males yield similar number of progeny. (d) Yellow has similar number of unfertilized oocytes compared to OP50. HT115, HB101, Red and Orange have significantly fewer unfertilized oocytes. Statistical comparisons by Tukey’s multiple comparison test. *, p<0.05; **, p<0.01; ***, p<0.001; ****, p<0.0001. (e) Volcano plot of differentially expressed reproduction genes on all bacterial foods compared to OP50. The number of significant genes have a p-value of <0.01. (f) Heat map of reproduction genes that are significant with a p-value <0.01 in at least one of the five bacterial diets, relative to the OP50 bacterial diet.

### Bacterial foods differentially alter organismal life expectancy and health

The quantity of diet ingested has been shown to influence lifespan across organisms^46,47^. Similarly, diet quality and composition can also influence healthspan, but the underlying mechanisms of healthspan improvement, and its relationship to diet, remain underdeveloped. As such, additional models to explore how diet can impact life- and healthspan are of great interest. To this end, we examined impact of each new food on organismal lifespan. Similar to previous studies^13^ animals raised on HB101 had a modest increase in mean lifespan, with the most significant impact on the last quartile of life (**Fig. 6a** and Supplementary Data 2). In contrast, worms raised on HT115, Red, and Yellow displayed significantly increased lifespan (**Figs. 6b-d** and Supplementary Data 2) while animals raised on Orange were short-lived (**Fig. 6e** and Supplementary Data 2). Taken together, our results show that Red, Yellow, and Orange represent three bacterial foods of sufficient nutritional quality to accelerate development, but differentially alter life expectancy in a food-dependent manner.

**Figure 6.**
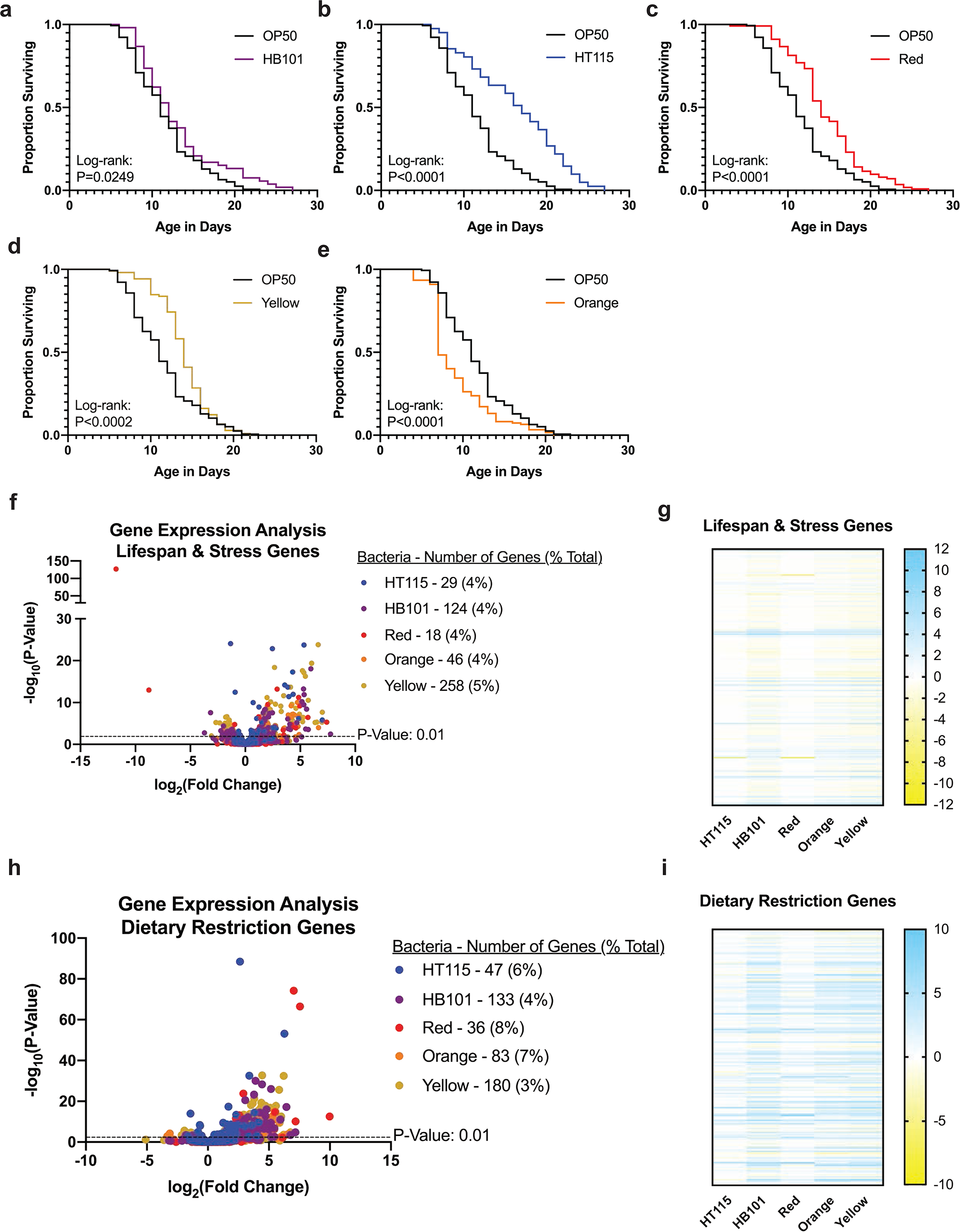
Lifespan of *C. elegans* on each bacterial diet. (a-e) Lifespan comparisons of OP50 versus each bacterial diet. HB101 (a) had a very small significant difference in lifespan compared to OP50. HT115 (b), Red (c) and Yellow (d) worms had increased lifespans and Orange (e) worms had a shorter lifespan. Lifespan comparisons between OP50-reared worms and the other diets made with Log-rank test. For lifespan quartile comparisons, refer to Supplementary Data 2. OP50 n=155, HT115 n=51, HB101 n=53, Red n=113, Orange n=122, Yellow n=105. (f) Volcano plot of differentially expressed genes related to lifespan and stress on all bacterial foods compared to OP50. The number of significant genes have a p-value of <0.01. (g) Heat map of lifespan and stress-related genes that are significant with a p-value <0.01 in at least one of the five bacterial diets. (h) Volcano plot of differentially expressed genes related to dietary restriction on all bacterial foods compared to OP50. The number of significant genes have a p-value of <0.01. (i) Heat map of dietary restriction-related genes that are significant with a p-value <0.01 in at least one of the five bacterial diets, relative to the OP50 bacterial diet.

We examined the transcriptional profiles of genes previously annotated to be involved in lifespan and discovered several interesting trends that may explain the lifespan-altering effects of each bacterial diet (**Figs. 6f-g**). Feeding the Yellow bacteria results in an increase in lifespan and, correspondingly, genes in the insulin-like signaling pathway (*daf-2*/Insulin receptor and *age-1*/PI3K) were downregulated, while DAF-16/FoxO target genes (Dao and Dod) were upregulated. Similarly, although less significantly, animals fed Red bacteria have increased expression of several genes regulated by DAF-16. Moreover, HT115, Red and Yellow all extend lifespan and share 11 genes with altered expression, all of which are upregulated. In previous studies, these 11 genes (*abu-1, abu-11, abu-14, gst-10, kat-1, lys-1, pqn-2, pqn-54, lpr-5, glf-1, bus-8*) lead to a shortened lifespan phenotype when the expression is reduced by RNAi^34^. Finally, feeding worms the Orange bacteria significantly decreased lifespan and although we did not identify any known longevity genes to display transcriptional changes, we found that *kgb-1, pha-4,* and *vhp-1* were all up regulated on this diet, while previous studies have observed extended lifespan when these genes are targeted by RNAi^34^. Because studies done in the past have shown that dietary restriction can influence lifespan^1^, we decided to also look at a subset of genes shown to differ between ad libitum and dietary restricted conditions ^48^. Interestingly, there was a similar proportion of genes in worms on each bacterial diet that were differentially expressed (**Figs. 6h-i**). Furthermore, Red and HT115 were more closely related to the OP50-raised worm genetic profile while HB101, Orange, and Yellow contained a higher number of genes that were both up and downregulated. Collectively, while longevity is a complex phenotype, feeding any of these bacterial diets can evoke transcriptional changes in genes associated with life expectancy.

Since lifespan can be uncoupled from healthspan^10,49^ we examined the impact of each bacterial diet on animal muscle function via thrashing as a surrogate for health (**Figs. 7a-e** and Supplementary Fig. 6). We measured thrashing rate at five specific timepoints that were selected for their significance in relation to other phenotypes: L4 larval development stage, based on the RNAseq data and fat staining analysis; day 1 of adulthood, based on the pharyngeal pumping assay; day 3 of adulthood, based on the presentation of the Asdf phenotype; day 8 of adulthood, due to 50% of the Orange population being dead at this time point; and day 11 of adulthood when 50% of the OP50 population had perished. Surprisingly, animals raised on HT115, HB101, Red, and Yellow had faster muscle movement while Orange were indistinguishable from OP50-fed animals at L4 stage (**Fig. 7a** and Supplementary Data 3) and at day 1 of adulthood (**Fig. 7b** and Supplementary Data 3). Although most day 3 animals had similar muscle function, animals fed HT115 and Orange had modestly slower movements (**Fig. 7c** and Supplementary Data 3). By day 8 of adulthood, Red-fed animals displayed significantly slower movements (**Fig. 7d** and Supplementary Data 3). Day 11 adults raised on HT115, HB101, Yellow, and Orange moved half as fast as they did at the end of development, similar to OP50-raised control animals, but Red-fed animals we significantly slower (~75% reduction) (**Fig. 7e** and Supplementary Data 3). Taken together, despite some bacteria enhancing early life muscle function, age-related decline remained similar, and was even enhanced in animals fed Red (**Fig. 7f**). It is notable that when examining the GO-term analysis of the 30 shared genes that are deregulated on all five of the bacterial foods, as compared to OP50, all 30 genes have been annotated as causing movement defects in *C. elegans* when the expression of that gene is altered^50–52^.

**Figure 7.**
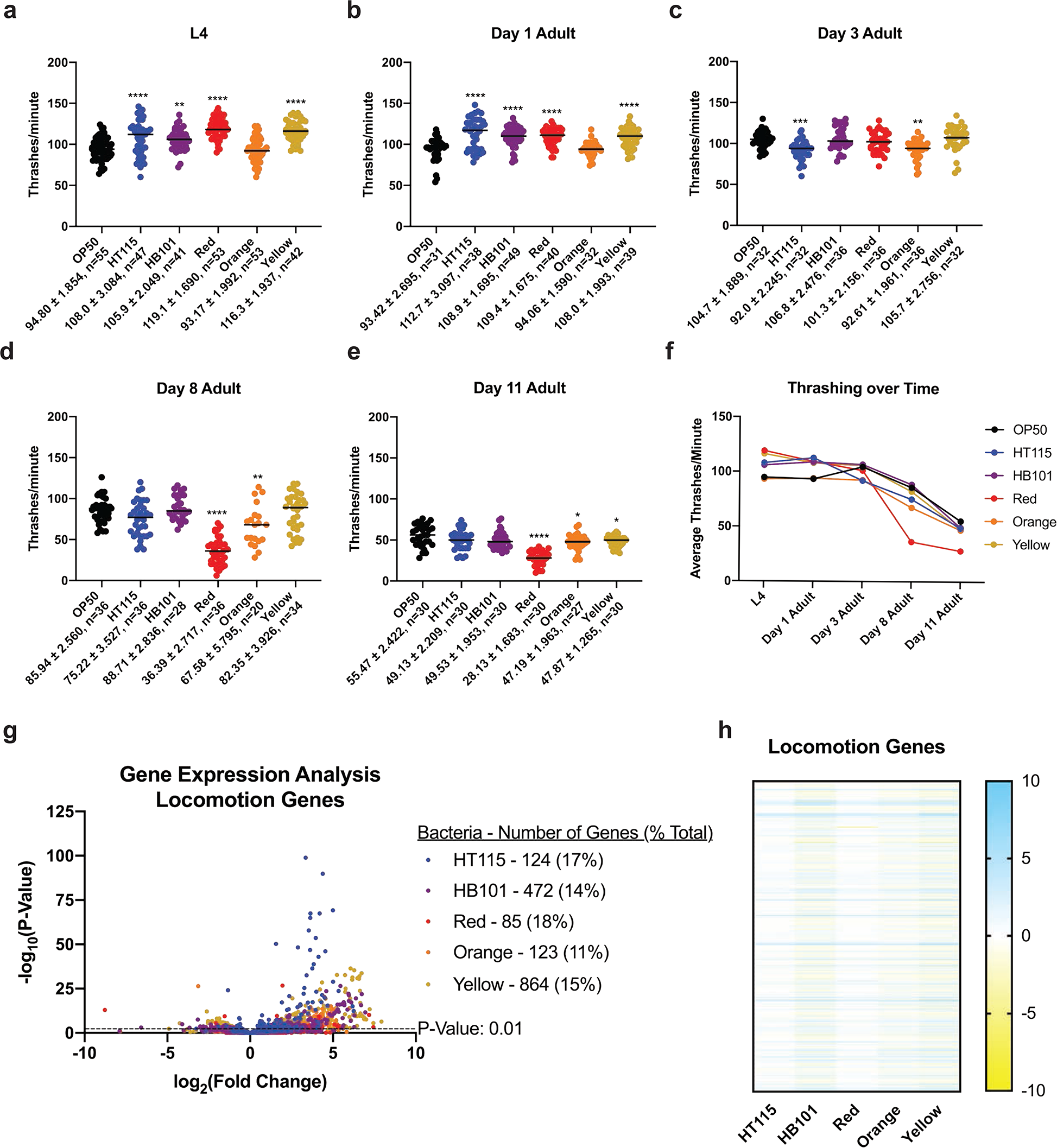
*C. elegans* have a food-dependent decline in thrashing with age. (a-e) The rate of thrashing in worms raised on each bacterial diet at different developmental stages. All comparisons were made to OP50-reared worms. (a) All foods except for Orange showed a significantly higher thrashing rate at the L4 stage. (b) All foods except for Orange showed a significantly higher thrashing rate at the day 1 adult stage. (c) HT115 and Orange thrashed significantly less than OP50 while the other bacterial diets were similar in thrashing rate at the day 3 adult stage. (d) Red and Orange had significantly lower thrashing rates compared to OP50 and all other bacterial foods at the day 8 adult stage. (e) Red, Orange, and Yellow had lower thrashing rates at day 11 adult when compared to OP50. Statistical comparisons by Tukey’s multiple comparison test. *, p<0.05; **, p<0.01; ***, p<0.001; ****, p<0.0001. (f) Average rate of thrashing over time on each bacterial diet. All studies performed in biological triplicate; refer to Supplementary Data 3 for n for each comparison. (g) Volcano plot of differentially expressed locomotion genes on all bacterial foods compared to OP50. The number of significant genes have a p-value of <0.01. (h) Heat map of locomotion genes that are significant with a p-value <0.01 in at least one of the five bacterial diets, relative to the OP50 bacterial diet.

### C. elegans are least often found dwelling on the Red bacteria when given the choice of other bacteria

Previously it has been shown that *C. elegans* display behaviors indicative of preference for bacterial diets that aid in better development, reproduction, and lifespan. Previous studies have suggested that dietary choices are made based on both the quality of food and the impact on the survival of the bacterial-feeding nematodes. Importantly, many animals select their food according to their environmental and dietary requirements ^53,54^. The overall trend of these past studies suggests *C. elegans* are choosing food sources aiding in higher fitness. Knowing that the six bacterial foods used in this study are capable of exerting diverse phenotypic changes (**Table 1**), we wanted to determine if foods we observed as more beneficial for longevity and healthspan would be what the worms were found feeding on when given multiple options. We set up a food choice assay to test this hypothesis with the use of OP50-reared worms (**Fig. 8a**). The food choice assay had the option of all six bacterial diets (**Fig. 8b**). When OP50-reared worms had the option to choose from any of the six bacteria, a trend was observed where the Red bacteria would be the only food source lacking dwelling worms. Taken together, this data suggests that *C. elegans* will choose any of the five bacterial foods over that of the Red bacteria, which is interesting since the Red bacteria promotes longevity.

**Table 1.**
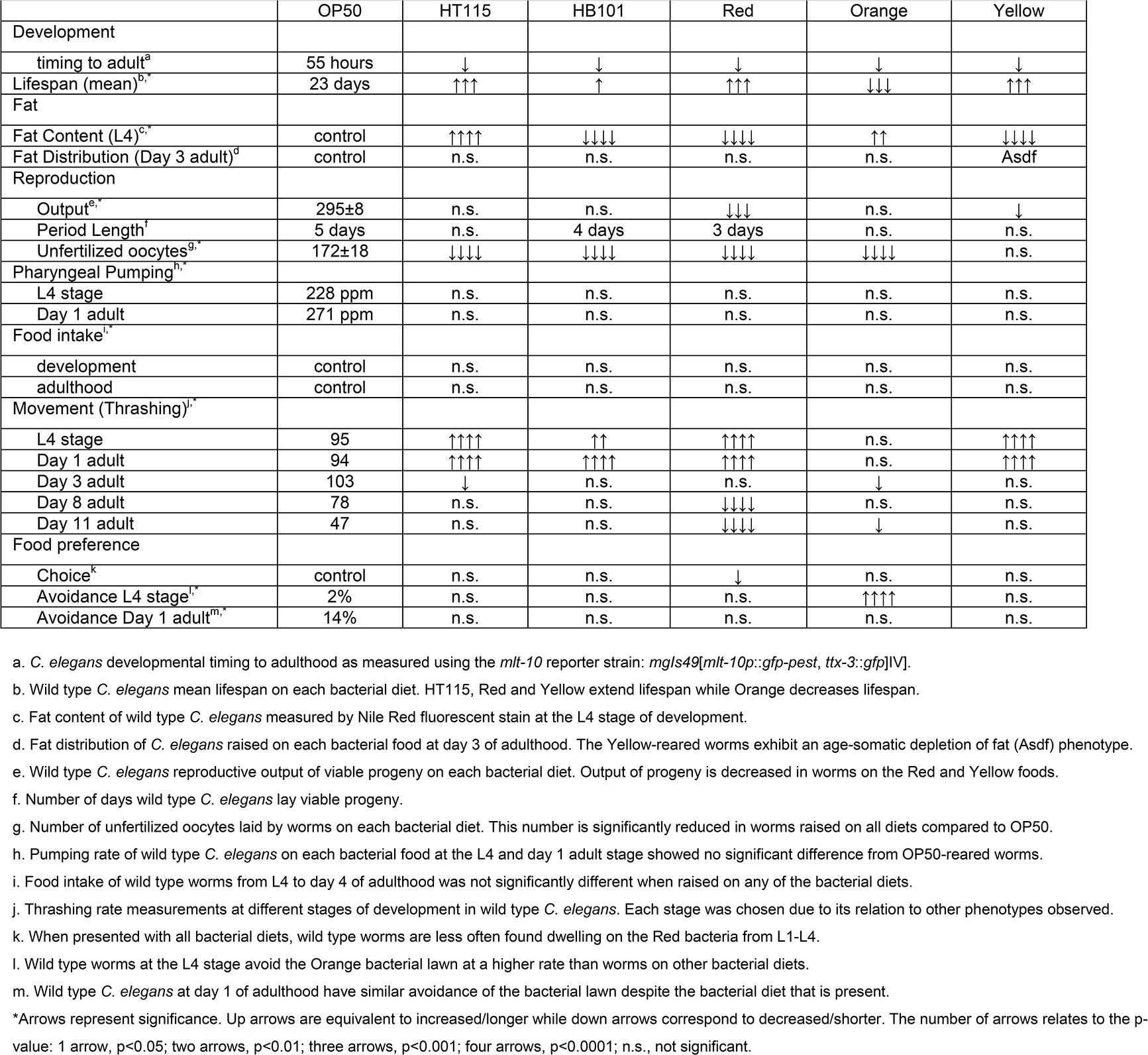
Bacterial diet exposure leads to changes in *C. elegans* physiology.

|  | OP50 | HT115 | HB101 | Red | Orange | Yellow |
| --- | --- | --- | --- | --- | --- | --- |
| Development |  |  |  |  |  |  |
| timing to adult<sup>a</sup> | 55 hours | ↓ | ↓ | ↓ | ↓ | ↓ |
| Lifespan (mean)<sup>b,*</sup> | 23 days | ↑↑↑ | ↑ | ↑↑↑ | ↓↓↓ | ↑↑↑ |
| Fat |  |  |  |  |  |  |
| Fat Content (L4)<sup>c,*</sup> | control | ↑↑↑↑ | ↓↓↓↓ | ↓↓↓↓ | ↑↑ | ↓↓↓↓ |
| Fat Distribution (Day 3 adult)<sup>d</sup> | control | n.s. | n.s. | n.s. | n.s. | Asdf |
| Reproduction |  |  |  |  |  |  |
| Output<sup>e,*</sup> | 295±8 | n.s. | n.s. | ↓↓↓ | n.s. | ↓ |
| Period Length<sup>f</sup> | 5 days | n.s. | 4 days | 3 days | n.s. | n.s. |
| Unfertilized oocytes<sup>g,*</sup> | 172±18 | ↓↓↓↓ | ↓↓↓↓ | ↓↓↓↓ | ↓↓↓↓ | n.s. |
| Pharyngeal Pumping<sup>h,*</sup> |  |  |  |  |  |  |
| L4 stage | 228 ppm | n.s. | n.s. | n.s. | n.s. | n.s. |
| Day 1 adult | 271 ppm | n.s. | n.s. | n.s. | n.s. | n.s. |
| Food intake<sup>i,*</sup> |  |  |  |  |  |  |
| development | control | n.s. | n.s. | n.s. | n.s. | n.s. |
| adulthood | control | n.s. | n.s. | n.s. | n.s. | n.s. |
| Movement (Thrashing)<sup>j,*</sup> |  |  |  |  |  |  |
| L4 stage | 95 | ↑↑↑↑ | ↑↑ | ↑↑↑↑ | n.s. | ↑↑↑↑ |
| Day 1 adult | 94 | ↑↑↑↑ | ↑↑↑↑ | ↑↑↑↑ | n.s. | ↑↑↑↑ |
| Day 3 adult | 103 | ↓ | n.s. | n.s. | ↓ | n.s. |
| Day 8 adult | 78 | n.s. | n.s. | ↓↓↓↓ | n.s. | n.s. |
| Day 11 adult | 47 | n.s. | n.s. | ↓↓↓↓ | ↓ | n.s. |
| Food preference |  |  |  |  |  |  |
| Choice<sup>k</sup> | control | n.s. | n.s. | ↓ | n.s. | n.s. |
| Avoidance L4 stage<sup>l,*</sup> | 2% | n.s. | n.s. | n.s. | ↑↑↑↑ | n.s. |
| Avoidance Day 1 adult<sup>m,*</sup> | 14% | n.s. | n.s. | n.s. | n.s. | n.s. |

a. *C. elegans* developmental timing to adulthood as measured using the *mlt-10* reporter strain: *mgIs49[mlt-10p::gfp-pest, ttx-3::gfp]IV*.
b. Wild type *C. elegans* mean lifespan on each bacterial diet. HT115, Red and Yellow extend lifespan while Orange decreases lifespan.
c. Fat content of wild type *C. elegans* measured by Nile Red fluorescent stain at the L4 stage of development.
d. Fat distribution of *C. elegans* raised on each bacterial food at day 3 of adulthood. The Yellow-reared worms exhibit an age-somatic depletion of fat (Asdf) phenotype.
e. Wild type *C. elegans* reproductive output of viable progeny on each bacterial diet. Output of progeny is decreased in worms on the Red and Yellow foods.
f. Number of days wild type *C. elegans* lay viable progeny.
g. Number of unfertilized oocytes laid by worms on each bacterial diet. This number is significantly reduced in worms raised on all diets compared to OP50.
h. Pumping rate of wild type *C. elegans* on each bacterial food at the L4 and day 1 adult stage showed no significant difference from OP50-reared worms.
i. Food intake of wild type worms from L4 to day 4 of adulthood was not significantly different when raised on any of the bacterial diets.
j. Thrashing rate measurements at different stages of development in wild type *C. elegans*. Each stage was chosen due to its relation to other phenotypes observed.
k. When presented with all bacterial diets, wild type worms are less often found dwelling on the Red bacteria from L1-L4.
l. Wild type worms at the L4 stage avoid the Orange bacterial lawn at a higher rate than worms on other bacterial diets.
m. Wild type *C. elegans* at day 1 of adulthood have similar avoidance of the bacterial lawn despite the bacterial diet that is present.
\*Arrows represent significance. Up arrows are equivalent to increased/longer while down arrows correspond to decreased/shorter. The number of arrows relates to the p-value: 1 arrow, p<0.05; two arrows, p<0.01; three arrows, p<0.001; four arrows, p<0.0001; n.s., not significant.

**Figure 8.**
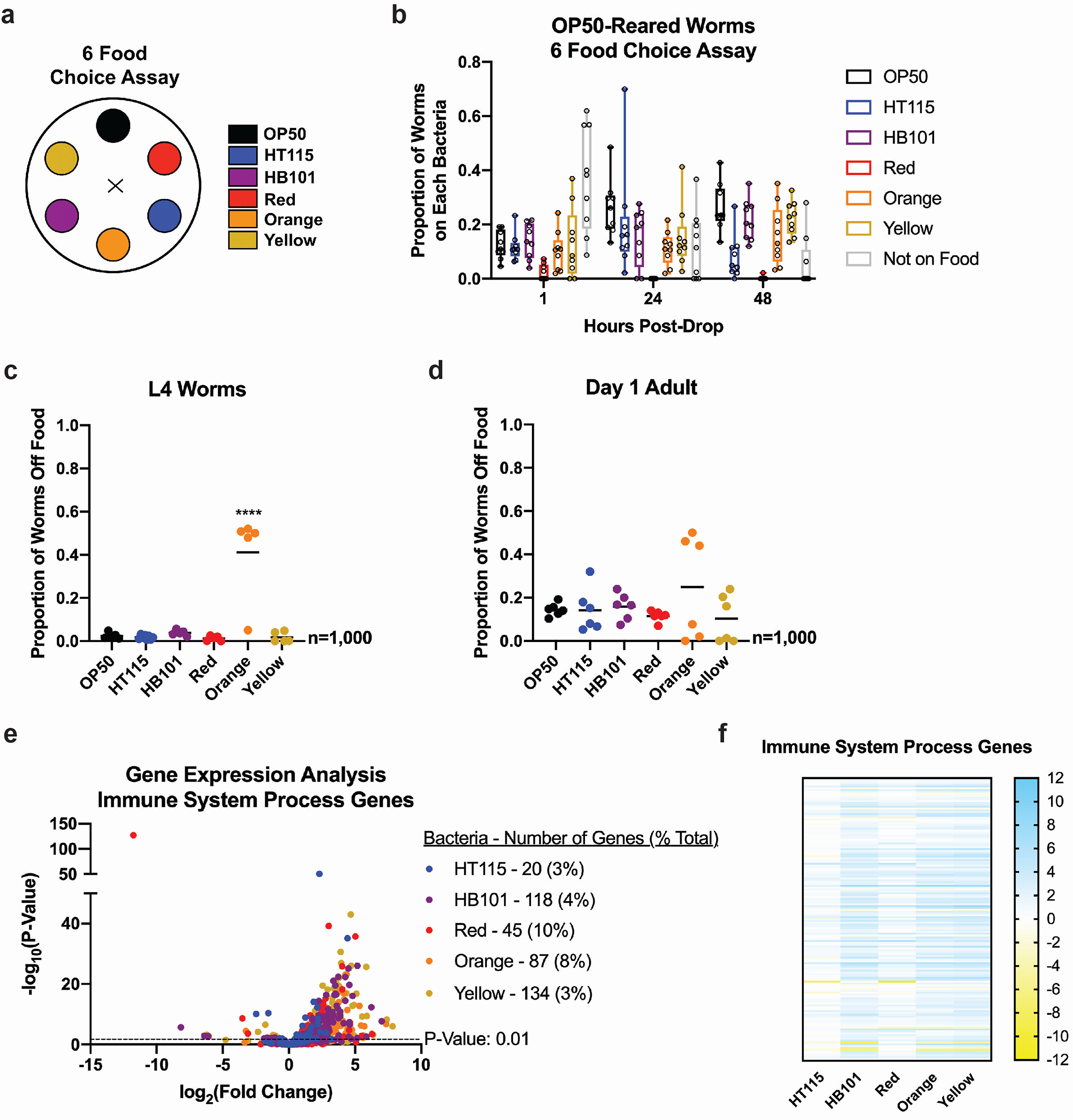
*C. elegans* food choice. (a) Schematic of the food choice assay. (b) Food choice of *C. elegans* raised on OP50 when presented with all six bacterial foods. Worms were found less often to be dwelling on the Red bacteria compared to the other bacterial diets. Each data point represents one plate of 50 worms (9 plates total; n=450 worms assessed). Each box represents 95% confidence intervals with the median value defined. (c) Proportion of worms on and off of the bacterial lawn at the L4 stage of development. n=200/replicate for a total of 5 replicates. (d) Proportion of worms on and off of the bacterial lawn at day 1 of adulthood. n=200/replicate for a total of 5 replicates. (e) Volcano plot of differentially expressed immune-related genes on all bacterial foods compared to OP50. The number of significant genes have a p-value of <0.01. (f) Heat map of immune-related genes that are significant with a p-value <0.01 in at least one of the five bacterial diets, relative to the OP50 bacterial diet.

The standard monoculture of OP50 in the laboratory environment is generally nonpathogenic and wild type worms typically remain on the bacterial lawn until all bacteria is eaten. In the case of pathogenic bacteria, *C. elegans* tend to initially eat the bacteria when encountered and then move off of the lawn in a span of a couple hours, a term commonly called lawn avoidance ^55,56^. In order to determine possible reasons for why *C. elegans* are less commonly found dwelling on the Red bacteria, we employed a lawn avoidance assay, which counts the number of worms residing on and off of a bacterial lawn at the L4 stage (**Fig. 8c**) and day 1 adult stage (**Fig. 8d**) of development. These assays revealed that the Red bacteria does not induce an avoidance response at either L4 or the day 1 adult stage. In fact, the proportion of worms found on the Red bacterial lawn is equivalent to the proportion of worms found on most of the other bacterial lawns. The one outlier to this is the Orange bacteria during the L4 stage, which has about forty percent of the animals residing off of the bacterial lawn. Knowing that pathogenic bacteria evoke an immune response, we asked if this was reflected in the RNA sequencing data. We examined a list of genes that are commonly found to be altered in worms when in the presence of pathogenic bacteria and found that the proportion of genes differentially expressed in worms on each bacterial diet ranged from 3-10% (**Fig. 8e**). When comparing the expression profiles of these immune response genes, we found that HT115 and Red bacteria resulted in similar gene expression responses while Orange and Yellow were closer to HB101 (**Fig. 8f**).

### Bacterial diet exposure at different timepoints in development alter lifespan trajectories

Previous studies have demonstrated that effects of calorie restriction (CR) can be realized even when initiated later in life^57,58^. Moreover, when fed ad libitum after a period of CR, mortality is shifted as if CR never occurred. With this model in mind, we asked if switching bacteria type, rather than abundance, could be wielded to alter lifespan outcomes. To address these questions, we switched growth conditions at major life stages (development, reproduction, post-reproduction) and measured lifespan (**Fig. 9a**). We used both Orange bacteria, which decreases lifespan, and Red bacteria, which increases lifespan, as the basis of our model. Remarkably, after assessing eight combinations of bacterial diet switching, we discovered that the bacteria fed during development (food 1) and reproductive period (food 2) contributed little to overall lifespan and the last bacteria exposed (food 3) exerted the most impact. In brief, when animals were exposed to the Red bacteria after experiencing the Orange bacteria, the normally shortened lifespan resulting from ingestion of Orange is suppressed (**Figs. 9b-d** and Supplementary Data 4). Conversely, exposure to Orange suppresses the normally extended lifespan linked to whole-life feeding of the Red bacteria (**Figs. 9e-g** and Supplementary Data 4). Intriguingly, when compared to animals raised on OP50, ingestion of Red bacteria post-reproductively, regardless of ingestion of Orange bacteria at any other life stage, results in a relatively normal lifespan; not shortened (Supplementary Fig. 7). Moreover, animals that eat the Red bacteria post-reproductively have an extended lifespan, which is further enhanced if the food is introduced after development. Taken together, our studies reveal that expanding the available foods for *C. elegans* is a powerful tool to study the impact of food on lifespan and healthspan. Future studies to integrate genetic analyses to define new gene-diet pairs^6,59^ and gene-environment interactions in general, will be of significant interest.

**Figure 9.**
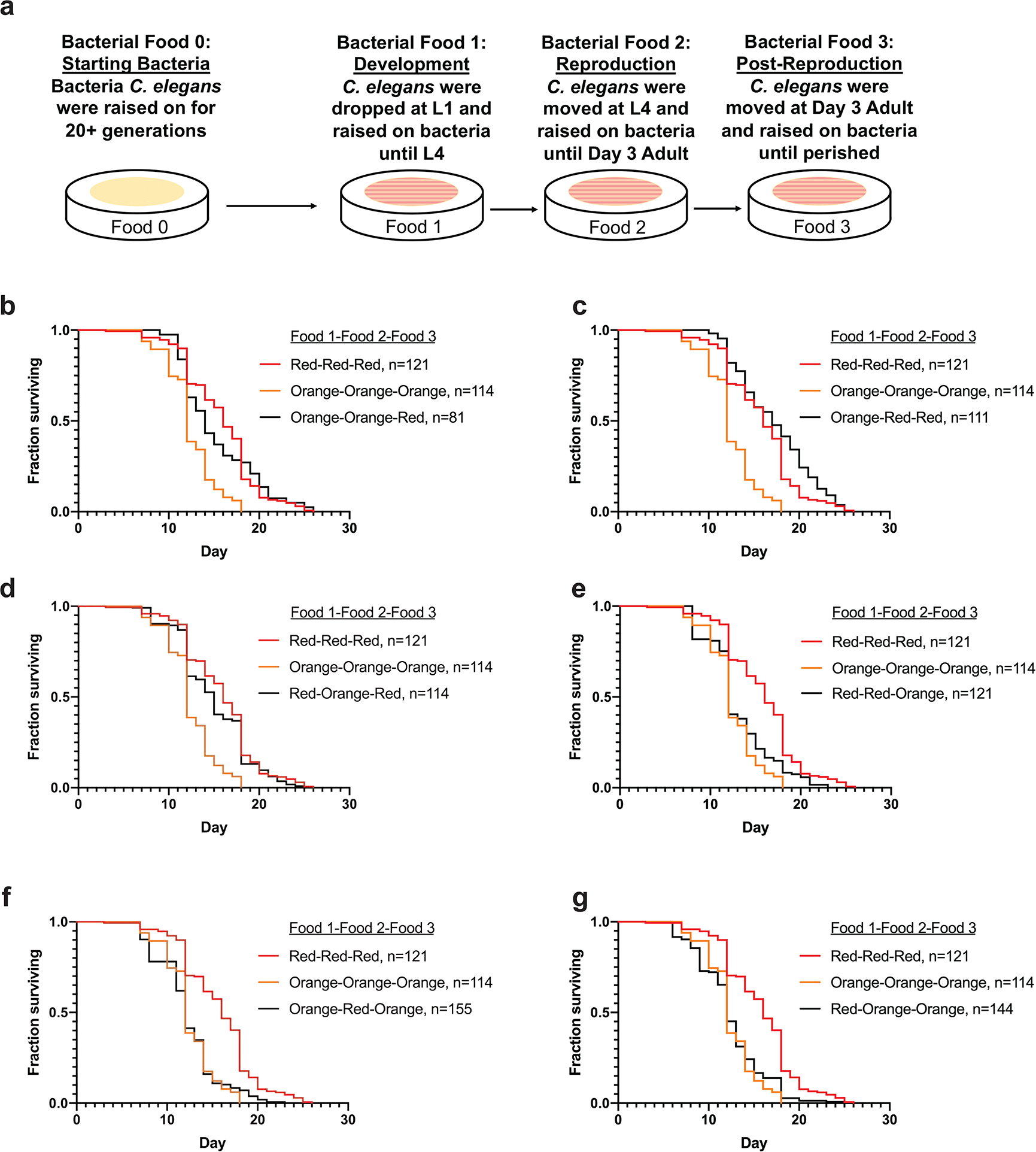
The introduction of the Red bacteria at the post-reproductive stage in *C. elegans* extends lifespan. (a) Schematic of the bacterial diet as a nutraceutical experiment. Food 0 was OP50 for all worms. After synchronization and allowing L1s to hatch overnight, L1s were dropped on bacterial food 1 and moved at L4 to bacterial food 2 before being moved to bacterial food 3 at day 3 of adulthood. Lifespans were then measured. (b-g) Lifespan curves of *C. elegans* on each of the different bacterial diet combinations. Lifespan comparisons between the bacterial diet combinations and Red-only, and Orange-only were made with Log-rank test; refer to Supplementary Data 4. *, p<0.05; **, p<0.01; ***, p<0.001; ****, p<0.0001.

## DISCUSSION

*C. elegans* is a well-established model to study diet and aging. Here we augment that model by introducing a comprehensive phenotypic analysis of *C. elegans* fed *E. coli* and three laboratory bacteria originally isolated from contaminated plates: *Methylobacterium*, *Xanthomonas*, and *Sphingomonas*. Interestingly, these microbes have been found in the normal *C. elegans* environments^16^ and when compared to the three most common *E. coli* strains OP50, HT115, HB101, our study is a critical tool to aid in our understanding of how these foods influence physiological and transcriptomic responses over the lifespan^11,16^. Previous works have identified how *C. elegans* react to the three standard *E. coli* foods provided in the lab^5,8,59–63^ and yet, to our knowledge, a “head-to-head" comparison of the age-related and healthspan-relevant outcomes that result from feeding these bacteria, is lacking. Because of this, we decided to investigate the effects of bacterial diet on both physiological attributes and transcriptional signatures of *C. elegans* raised on bacterial species for thirty generations, to avoid acute stress responses to the food.

Recent studies have shown that bacterial diet can alter transcriptional responses in *C. elegans*^64,65^. Our work supports this observation with three new menu options for *C. elegans* culture. Although each of these foods evokes a unique transcriptional signature (**Fig. 2**), several specific classes of genes are shared among multiple, or even all of the bacterial diets. Our RNAseq analysis was limited to gene expression changes observed as animals enter adulthood, prior to reproduction. Given the extent of changes in reproductive capacity, health (movement and fat), and aging, future work to examine how gene expression is altered on each food over the lifespan will be of great interest. Moreover, a fine-tuned analysis of transcription at each development stage will be informative based on our observation that different foods can accelerate the transition of animals across specific developmental stages. Nevertheless, it is clear that the standard laboratory *E. coli* strains and our three new bacterial foods induce a transcriptional response of phenotypically relevant genes. Knowing that even different strains of *E. coli* can differentially impact physiology and gene expression, we acknowledge that our findings may not be generalizable for all *Methylobacterium*, *Xanthomonas*, and *Sphingomonas* and are potentially strain dependent.

Bacteria serve as a live food source for *C. elegans* both in their natural environment and laboratory setting. Prior studies have shown animals pumping at similar rates in the presence of bacteria they can and cannot eat^12,66^. The rates of pharyngeal pumping were not significantly different on any of the bacteria in L4 stage and day 1 adult animals (**Fig. 4o** and Supplementary Fig. 4a). Because pharyngeal pumping may not be the best way to measure overall consumption of food, we employed a food intake assay to measure the quantity of food ingested by worms fed each bacterial diet. These data demonstrated that worms were eating at similar rates on the bacterial diets between the L4 larva and day 4 adult stages (**Fig. 4p**). We also found that when examining the bacterial load in day 1 adults, it appears that all bacterial diets are able to be broken down and digested by the worms due to the lack of colony growth on LB plates after the lysis of individual worms (Supplementary Fig. 4h). In support of this data, we also observed faster developmental timing (**Fig. 3**) and large-sized broods (**Fig. 5a**), indicating that each bacterium is providing sufficient nutrition to the worms feeding on them. Nevertheless, based on the similarities of animals eating Red and animals undergoing dietary restriction, it remains possible that this bacterium allows ad libitum ingestion with the physiological benefits of reduced eating, which is further supported by the Red bacteria causing worms to have the highest proportion of dietary restriction-related genes deregulated in comparison to the other bacterial diets (**Figs. 6h-i**).

Clearly, changing food sources can potently impact multiple phenotypic attributes and several of these aging-relevant phenotypes are interrelated. For example, reproduction, fat, and stress resistance are intrinsically tied to overall life expectancy^9^. In support of this model, the Red and Yellow bacteria, which reduce overall reproduction (**Fig. 5a**), indeed increase lifespan (**Fig. 6**). However, our study reveals that this relationship is more complex as these two foods have different effects on lipid storage and Yellow evokes an age-dependent somatic depletion of fat (Asdf) response (**Fig. 4**), which is a phenotype observed in animals exposed to pathogens^36,37^. It may be that *C. elegans* perceive the ingested Yellow bacteria as a pathogen, however the increased lifespan that follows on this diet, suggests this food is health-promoting. The Yellow bacteria also does not produce a lawn avoidance phenotype at L4 or day 1 of adulthood (**Figs. 8c-d**), which is a common phenotype that occurs within hours of introduction of worms to pathogenic bacteria. Additionally, these data are supported by the RNAseq data analysis of immune system-related genes, which shows that the Yellow bacteria induces a proportionally smaller number of gene expression changes compared to the other bacterial diets used in this study (**Figs. 8e-f**). Lastly, genetic and environmental mechanisms that delay developmental timing have been tied to increased longevity in adulthood^40,67,68^. Each of the bacterial diets we tested results in faster development into a reproductive adult (**Fig. 3**), but only HB101, HT115, Red, and Yellow increase lifespan, while Orange decreases lifespan. Taken together our study reveals that each bacterial food can exert a specific life history changing response, but also questions previously established models of aging.

Many aspects of the bacterial communities, besides that of their nutritional composition, have been shown to influence multiple attributes in *C. elegans,* including food preference, feeding rates, brood size, and lifespan^69,70^. We decided to explore the food preference aspect by carrying out a food choice assay that contained all six of our bacterial diets. We hypothesized that bacterial diets shown to extend lifespan, like Red and Yellow (**Figs. 6c-d**), would have higher levels of dwelling worms while the Orange bacteria would have fewer dwelling worms due to the shortened lifespan (**Fig. 6e**). Interestingly, in the food choice assay, we saw that worms are less often found on the Red bacteria and more often found on the other bacteria. Finding worms more often on both Yellow and Orange was somewhat surprising due to the Asdf phenotype at day 3 of adulthood in Yellow-raised worms (**Figs. 4m-n**) and the shortened lifespan on Orange (**Fig. 6e**). Both of these phenotypes are consistent with phenotypes when worms are exposed to pathogens. However, the pathogenic effect of specific bacteria is often associated with intestinal colonization ^71–73^, which we did not observe (Supplementary Fig. 4h). Moreover, the increased lifespan of animals fed the Yellow bacteria disagrees with the idea that the Yellow bacteria is pathogenic. Similarly, the lack of significant bacterial lawn avoidance at day 1 of adulthood when animals are reared on Orange bacteria suggests the Orange diet isn’t perceived as detrimental. Nevertheless, these results motivate future experimentation to understand why worms are found less frequently on a longevity-promoting bacterium and more frequently on a bacterium that decreases lifespan.

Inspired by previously described dietary interventions that are capable of extending lifespan in multiple species^47,74,75^, we wanted to investigate whether, instead of keeping animals on one bacterial diet their entire life, altering food exposure at critical life stages could affect overall lifespan. We asked whether these foods could alter lifespan without any previous generational exposure. Strikingly, acute exposure to these bacterial foods was capable of altering lifespan (**Fig. 9**), but intriguingly the magnitude of the response differed from animals that have been on these foods for 30+ generations (**Fig. 6**). Furthermore, our study revealed that the bacteria fed during the post-reproductive period exerted the strongest influence over the lifespan of the cohort. This result, however, is potentially confounded by the amount of time (longitudinally) that is spent on each bacteria. Regardless, our study reveals that any potential early-life (development and reproductive span) exposure to foods that normally shorten lifespan can be mitigated by eating a lifespan-promoting food option. This result is reminiscent of previous studies showing the mortality rate of dietary restricted (DR)-treated animals is accelerated when switched to an ad libitum diet while mortality rate is reduced when ad libitum-fed animals are switched to a DR diet^76^. In our case, calories are perhaps not different on Red or Orange, but rather the overall nutritional composition of the bacteria is different. Given that in our model animals are chronically exposed to as much food as they can eat throughout their life, we posit that if a similar human diet were discovered that this treatment protocol would be more accessible as food restriction is difficult, socially and psychologically^77–79^.

Ultimately, this study reveals the impact that diet can have on both physiology and the transcriptomics of animals that thrive on them. These alterations caused by differential bacterial diet exposure can not only be seen at the surface level in the diverse phenotypes that present themselves, but also at the genetic level, causing fluctuations in gene expression important for multiple physiological processes including development, fat content, reproduction, healthspan, and lifespan (**Table 1**). These discoveries in the worm can aid in understanding how dietary exposure influences different phenotypes and continuing work in the effects of bacterial diet on overall health and aging will unequivocally contribute to more personalized diets to promote healthier and longer lives in individuals. Taken together, our study expands the menu of bacterial diets available to researchers in the laboratory, identifies bacteria with the ability to drive unique physiological outcomes, and provides a food quality approach to better understand the complexities of gene-diet interactions for health over the lifespan.

## Supporting information

Supplemental materia

## ACKNOWLEDGEMENTS

H. Dalton for early work on this project including isolation of new bacterial contaminants, C-A Yen for experimental assistance, J. Gonzalez for technical assistance, A. Hammerquist for critical reading of the manuscript, A. Frand for the strain GR1395 (*mgIs49*[*mlt-10p*::*gfp-pest*, *ttx-3*::*gfp*]IV]), and members of the Curran lab for thoughtful suggestions.

## AUTHOR CONTRIBUTIONS

S.P.C. designed the study; N.L.S. performed the experiments; N.L.S. and S.P.C. analyzed data. S.P.C. wrote the draft manuscript, and N.L.S. and S.P.C. revised the final manuscript. This work was funded by the NIH R01GM109028 and R01AG058610 to S.P.C. and T32AG052374 and T32GM118289 to N.L.S.

## METHODS

### *C. elegans* strains and maintenance

*C. elegans* were raised on 6 cm nematode growth media (NGM) plates supplemented with streptomycin and seeded with each bacterial diet. For experiments, nematode growth media plates without streptomycin were seeded with each bacterium at the optical density of 0.8 A_600_. All worm strains were grown at 20°C and unstarved for at least three generations before being used. Strains used in this study were N2 Bristol (wild type) and GR1395 (*mgIs49*[*mlt-10p*::*gfp-pest*, *ttx-3*::*gfp*]IV]). Some strains were provided by the CGC, which is funded by NIH Office of Research Infrastructure Programs (P40 OD010440).

*E. coli* strains used: OP50, HT115(DE3), HB101. Red, Yellow, and Orange bacteria were isolated from stock plates in the laboratory and selected for with antibiotics before inoculating. Red, Orange, and Yellow were sequenced using the 16S primer pair 337F (GACTCCTACGGGAGGCWGCAG) and 805R (GACTACCAGGGTATCTAATC) and identified using the blastn suite on the NCBI website.

### Bacterial Growth Curves

Three test tubes with 5mL of LB without antibiotics were inoculated with each bacterium before being place in 37°C (except for Yellow, which was placed at 26°C). Optical density measurements were taken in duplicate every hour for 12 hours. The final measurement was taken 24 hours later. Optical density curves were made by averaging the readings together after performing the experiment in biological triplicate.

The antibiotic versus no antibiotic growth curves were conducted in a similar manner. Three test tubes with 5mL of LB with and without appropriate antibiotics (OP50/HB101 with streptomycin and HT115/Red/Orange/Yellow with ampicillin) were inoculated with each bacterium before being place in 37°C (except for Yellow, which was placed at 26°C). Optical density measurements were taken in duplicate at 12 hours and 24 hours after inoculation. Optical density curves were made by averaging the readings together after performing the experiment in biological triplicate.

### Metabolite Kits

Bacteria was grown overnight in LB liquid culture with corresponding antibiotics. The next day, cultures were collected at the log phase of growth, spun down, and bacteria was washed in water three times before being spun down and frozen at −80°C until use. Red bacteria samples were seeded onto LB plates and then allowed to grow overnight before scraping off and collecting. Bacteria were homogenized based on metabolite kit instructions and corresponding metabolites were measured via Bio Vison kit instructions. Both glycogen and glucose measurements were obtained from the Glycogen Colorimetric/Fluorometric Assay Kit (K646) and the triglyceride and glycerol measurements were obtained from the Triglyceride Quantification Colorimetric/Fluorometric Kit (K622).

### Bomb calorimetry

Bacteria was grown overnight in liquid culture of LB with corresponding antibiotics. The next day, bacteria were collected at the log phase, seeded onto LB plates at 0.8 optical density, and allowed to grow overnight. Bacteria was then collected with autoclaved water from the seeded plates and spun down to remove excess water. Samples were frozen at −80°C until being sent in triplicate to the Department of Nutrition Sciences at the University of Alabama at Birmingham for sample drying and bomb calorimetry. Water content measurements were also obtained after measuring wet weight and dry weight of the samples.

### RNA-sequencing

Wild type worms grown on each food for at least 30 generations were egg prepped and eggs were allowed to hatch overnight for a synchronous L1 population. The next day, L1s were dropped on NGM plates seeded with 0.8 OD of each bacterial diet. 48 hours post drop, L4 animals were washed 3 times with M9 buffer and frozen in TRI reagent at −80°C until use. Animals were homogenized and RNA extraction was performed via the Zymo Direct-zol RNA Miniprep kit (Cat. #R2052). Qubit™ RNA BR Assay Kit was used to determine RNA concentration. The RNA samples were sequenced and read counts were reported by Novogene. Read counts were then used for differential expression (DE) analysis using the R package DESeq2 created using R version 3.5.2. Statistically significant genes were chosen based on the adjust p-values that were calculated with the DESeq2 package. Genes were selected if their p-value<0.01.

### Microscopy

All images in this study were acquired using ZEN software and Zeiss Axio Imager. Worm area comparisons were imaged at 10x magnification (L2-L3 and L4 stage worms) and 5x magnification (day 1 adult and day 2 adult) with the DIC filter. Worm areas were measured in ImageJ using the polygon tool. For GFP reporter strains, worms were mounted in M9 with 10mM levamisole and imaged with DIC and GFP filters. For staining of bacteria, Biotium BactoView^TM^ Live Fluorescent Stain was used to visualize the DNA inside the cells.

### Developmental timing by *mlt-10::gfp*

GR1395 worms grown on each bacterial food for at least 20 generations were egg prepped and eggs were allowed to hatch overnight for a synchronous L1 population. One 24-well plate per food source, each containing a single L1 worm, were visualized by fluorescence microscopy every hour for 55 hours. Each hour worms were scored as green (molting) or non-green (not molting). Wells without worms, wells with two worms and worms that crawled to the side of the plate were censored.

### Lifespan analysis

Worm strains on each bacterial diet were egg prepped to generate a synchronous L1 population and dropped on the corresponding food. Worms were kept at 20°C and moved every other day during progeny output (day 2, day 4, day 6, day 8). Every diet was moved at the same time. Worms were scored daily for survival by gentle prodding with a platinum wire. Animals that burst or crawled to the side of the plate were censored and discarded from this assay.

### Reproduction assay

Wild type worms grown on each bacteria for at least 30 generations were egg prepped and eggs were allowed to hatch overnight for a synchronous L1 population. The next day, L1s were dropped on NGM plates seeded with each bacterial diet (OD 0.8). L4s hermaphrodites were singled 48 hours later onto individual plates and moved every 12 hours until egg laying ceased. Progeny were counted 48 hours after the singled hermaphrodite was moved to another plate. Progeny were removed from the plate during counting in order to ensure accuracy. Unfertilized oocytes were counted 24 hours after the singled hermaphrodite was moved to another plate.

### Mated reproduction

Male wild type populations were raised for at least 10 generations before mating to wild type hermaphrodite populations grown for 30+ generations on each bacteria. Wild type males raised on Red and OP50 were egg prepped at the same time as wild type hermaphrodite populations raised on Red and OP50 and allowed to hatch overnight in M9 to create synchronous L1 populations. The next day, L1s were dropped onto each corresponding diet and allowed to grow for 48 hours into L4s. L4 hermaphrodites were singled onto a plate with 30mL of OP50 or Red bacteria seeded in the center of the NGM plate. A single male was placed with each hermaphrodite and allowed to mate for 24 hours before the hermaphrodite was moved to a new plate and allowed to egg lay. Hermaphrodites were moved every day to a new plate until egg laying ceases. Progeny were counted 48 hours after moving the hermaphrodite and plates were checked for ~50% males to ensure the hermaphrodite was mated. Progeny were removed from the plate during counting in order to ensure accuracy.

### Nile Red Staining

Worms were grown on each bacterial diet for 48 or 72 hours, to the L4 stage or day 1 adult stage respectively and washed with 1x phosphate-buffered saline with Tween detergent (PBST) wash buffer. Worms were then rocked for 3 minutes in 40% isopropyl alcohol before being pelleted and treated with Nile Red in 40% isopropyl alcohol for 2 hours. Worms were pelleted after 2 hours and washed in PBST for 30 minutes before being imaged at 10x magnification (L4) or 5x magnification (day 1 adult) with DIC and GFP filters on the Zeiss Axio Imager. Fluorescence is measured via corrected total cell fluorescence (CTCF) via ImageJ and Microsoft Excel. CTCF = Worm Integrated Density-(Area of selected cell X Mean fluorescence of background readings) and normalized to the control of OP50-reared worms.

### Oil Red O Staining

Worms were grown on each bacterial diet for 48 or 120 hours, to the L4 stage or day 3 adult stage respectively and washed with PBST. Worms were then rocked for 3 minutes in 40% isopropyl alcohol before being pelleted and treated with Oil Red O in diH2O for 2 hours. Worms were pelleted after 2 hours and washed in PBST for 30 minutes before being imaged at 10x magnification (L4) or 5x magnification (day 3 adult) with the DIC filter on the Zeiss Axio Imager Erc color camera.

### Asdf Quantification

ORO-stained worms were placed on glass slides and a coverslip was placed over the sample. Worms were scored, and images were taken using a Zeiss microscope at 10× magnification. Fat levels of worms were placed into 3 categories: non-Asdf, intermediate, and Asdf. Non-Asdf worms display no loss of fat and are stained a dark red throughout most of the body (somatic and germ cells). Intermediate worms display significant fat loss from the somatic tissues, with portions of the intestine being clear, but ORO-stained fat deposits are still visible (somatic < germ cells). Asdf worms have had most, if not all, observable somatic fat deposits depleted (germ cells only).

### Pharyngeal pumping assays

Wild type worms grown on each bacterial diet for at least 30 generations were egg prepped and eggs were allowed to hatch overnight for a synchronous L1 population. The next day, L1s were dropped on NGM plates seeded with a bacterial diet (OP50, HT115, HB101, Red, Orange or Yellow). Before imaging, worms were singled onto plates seeded with a bacterial diet 1-2 hours before videoing pumping. For L4 pumping analysis, 10-12 worms were imaged 48 hours later via Movie Recorder at 8ms exposure using the ZEN 2 software at 10X magnification (Zeiss Axio Imager). For day 1 adult pumping analysis, 10-12 worms were imaged 72 hours after dropping L1s.

### Food intake

Food intake experiments were adapted from Gomez-Amaro et *al.* 2015 ^80^. Food intake was assessed in NGM liquid media without streptomycin in flat-bottom, optically clear 96-well plates with 150 mL total volume. Plates contained 10-40 worms per well. Bacteria was grown overnight in liquid culture of LB with corresponding antibiotics. The next day, bacteria were collected at the log phase, seeded onto NGM plates at 0.8 optical density, and allowed to grow overnight. Bacteria was then collected with autoclaved water from the seeded plates and spun down to wash and resuspended in water. Age-synchronized nematodes were seeded as L1 larvae and grown at 20°. Plates were sealed with parafilm to prevent evaporation and rocked continuously to prevent drowning of the nematodes. 5-fluoro-2’-deoxyuridine (FUDR) was added 48 hours after seeding at a final concentration of 0.12 mM. OD_600_ of each well was measured using a plate reader every 24 hours starting at L4 stage and ending at Day 4 of adulthood (144 hours after dropping L1s). The fraction of animals alive was scored microscopically at Day 4 of adulthood. Food intake per worms was calculated as bacterial clearance divided by worm number in well. Measurements were then normalized to the L4 to Day 1 Adult clearance rate for each bacterial diet.

### M9 thrashing assays

Wild type worms grown on each bacterial diet for at least 30 generations were egg prepped and eggs were allowed to hatch overnight for a synchronous L1 population. The next day, L1s were dropped on NGM plates seeded with each bacterial diet (OD 0.8). Worms were grown on each bacterial diet until the L4, day 1 adult, day 3 adult, day 8 adult or day 11 adult stage. Worms were then moved to an unseeded NGM plate to remove bacteria from the cuticle of the worms. After an hour, worms were washed with M9 and dropped in 5mL M9 drops onto a fresh NGM plate. After one minute, 10-12 worms were imaged via Movie Recorder at 50ms exposure using the ZEN 2 software (Zeiss Axio Imager).

### Food choice assays

Bacteria was grown overnight in liquid culture of LB with corresponding antibiotics. The next day, bacteria were collected at the log phase, 30 mL of each bacterium was seeded onto NGM plates with no antibiotics at 0.8 optical density, and allowed to grow overnight. In the 6 food choice assays, all bacteria were seeded 2 cm from the center point on a 6 cm plate. For the 6 food choice assay, seeding order was OP50, Red, HT115, Orange, HB101, and then Yellow. Once food choice assay plates were seeded and allowed to grow overnight, OP50-reared worms were egg prepped and eggs were allowed to hatch overnight for a synchronous L1 population. The next day, L1s were dropped into the center of the NGM plate and counted. The plate was then checked at 1, 24 and 48 hours to observe the location of worms. If worms were found on the bacterial lawn, then those worms were counted as on that food. Worms found outside bacterial lawns were counted as not on food. The proportion of worms found on each food or off of food was then calculated and graphed. Each assay was done in biological triplicate with technical triplicates for a total of nine plates.

### Bacterial load assay

Bacteria was grown overnight in liquid culture of LB with corresponding antibiotics. The next day, bacteria were collected at the log phase, seeded onto NGM plates at 0.8 optical density, and allowed to grow overnight. Worms reared on each bacterial food were egg prepped and eggs were allowed to hatch overnight for a synchronous L1 population. The next day, L1s were dropped onto NGM plates with each bacterial diet. Worms were allowed to grow until the day 1 adult stage (72 hours post-drop). Worms were then washed with water and then dropped on an unseeded NGM plate. Worms were allowed to crawl for an hour before washing again and dropping onto a new unseeded NGM plate. Worms were allowed to crawl for an hour before lysing 24 individual worms in 10 mL of worm lysis buffer. 5 mL of the supernatant was then seeded onto LB plates and allowed to grow 48 hours at 37°. Plates were checked at 24 and 48 hours for growth. This was done in duplicate for a total of 48 individual worms per bacterial diet.

### Lawn avoidance assay

Bacteria was grown overnight in liquid culture of LB with corresponding antibiotics. The next day, bacteria were collected at the log phase, seeded onto NGM plates at 0.8 optical density, and allowed to grow overnight. Worms reared on each bacterial food were egg prepped and eggs were allowed to hatch overnight for a synchronous L1 population. The next day, L1s were dropped onto NGM plates with each bacterial diet and counted. Plates were checked 48 hours later at the L4 stage and the number of worms on and off food were counted. This was repeated at the day 1 adult stage 24 hours later as well. The proportion of worms was calculated and graphed using Graphpad Prism 8.

### Statistics and Reproducibility

Data are presented as mean ± SEM. Comparisons and significance were analyzed in Graphpad Prism 8. Comparisons between more than two groups were done using ANOVA. For multiple comparisons, Tukey’s multiple comparison test was used and p-values are *p<0.05 **p<0.01 *** p<0.001 ****<0.0001. Lifespan comparisons were done with Log-rank test. Sample size and replicate number for each experiment can be found in figures and corresponding figure legends. This information is also in the experimental methods. Exact values for graphs found in the main figures can be found in Supplementary Data 5.

### DATA AVAILABILITY

RNA-sequencing data are deposited into the Gene Expression Omnibus (accession no. GSE152794). All other relevant data is available upon request from the corresponding author.

