## Supplemental materia for "Bacterial diets differentially alter lifespan and healthspan trajectories in *C. elegans*"

**SUPPLEMENTAL FIGURES**

**a**

| Name | Bacteria | Doubling Time | Antibiotic Sensitivity | Growth Conditions for Experiments in Liquid Culture | Growth Conditions on Plate |
|---|---|---|---|---|---|
| OP50 | *Escherichia coli* | 2 Hours | Ampicillin | LB + Streptomycin at 37C | LB + Streptomycin at 37C |
| HT115 | *Escherichia coli* | 1 Hour | Streptomycin | LB + Ampicillin at 37C | LB + Ampicillin at 37C |
| HB101 | *Escherichia coli* | 3 Hours | Ampicillin | LB + Streptomycin at 37C | LB + Streptomycin at 37C |
| Red | *Methylobacterium* | 23 Hours | Streptomycin | LB + Ampicillin at 37C | LB + Ampicillin at 30C |
| Orange | *Xanthomonas* | 5 Hours | Streptomycin | LB + Ampicillin at 37C | LB + Kanamycin at 37C |
| Yellow | *Sphingomonas* | 7 Hours | Streptomycin | LB + Ampicillin at 26C | LB + Ampicillin at 37C |

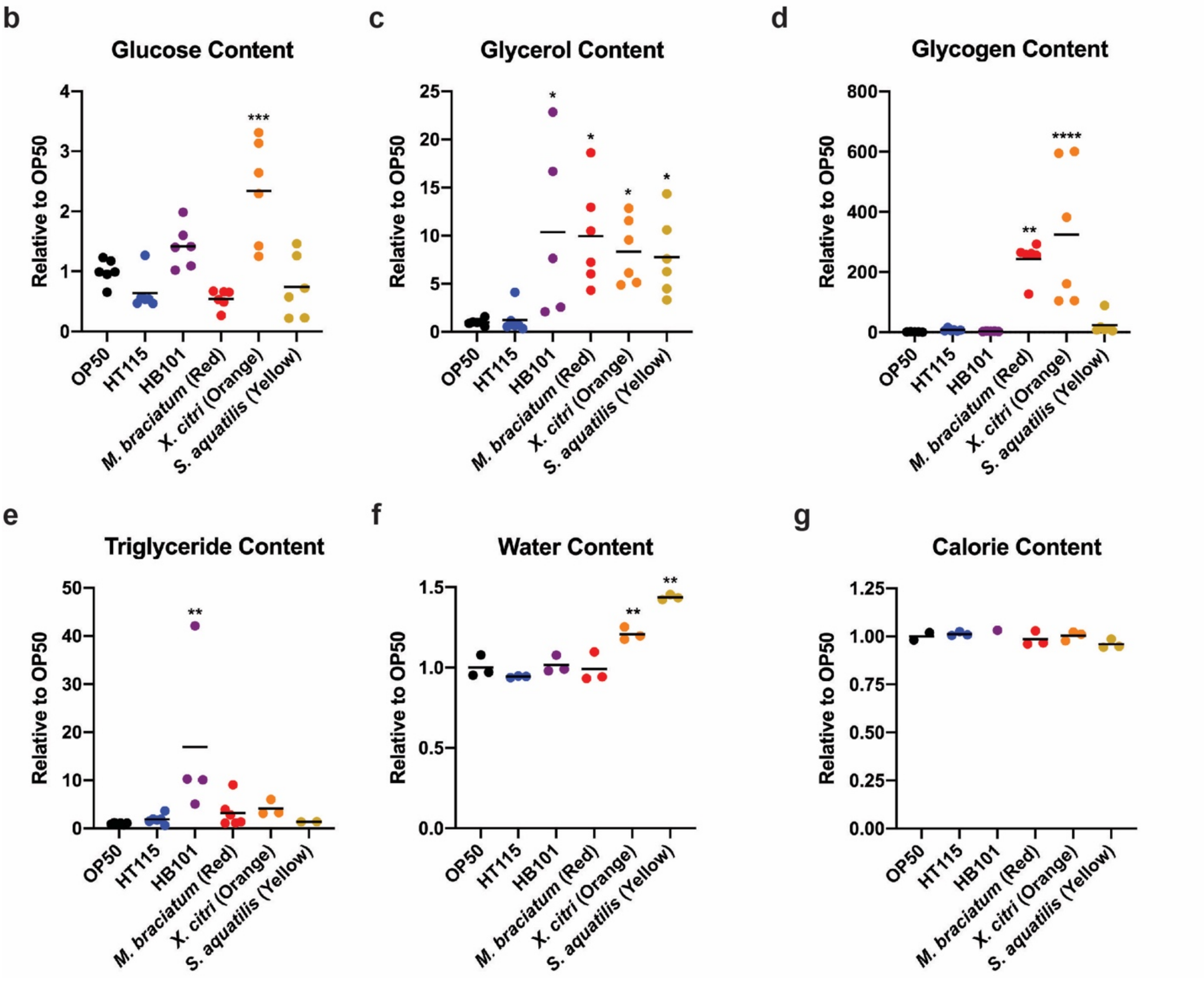


**Supplemental Figure 1. Bacterial growth conditions and metabolite concentrations.** (a) Table with optimal growth conditions for each bacterial diet. (b-g) Metabolite concentrations in each bacterium relative to OP50. The different metabolites measured were glucose (b), glycerol (c), glycogen (d), triglycerides (e) and water content (f). Statistical comparisons by Tukey's multiple comparison test. *, $p<0.05$; **, $p<0.01$; ***, $p<0.001$; ****, $p<0.0001$. All studies were performed in biological triplicate. (g) Calorie content of all bacteria were similar.

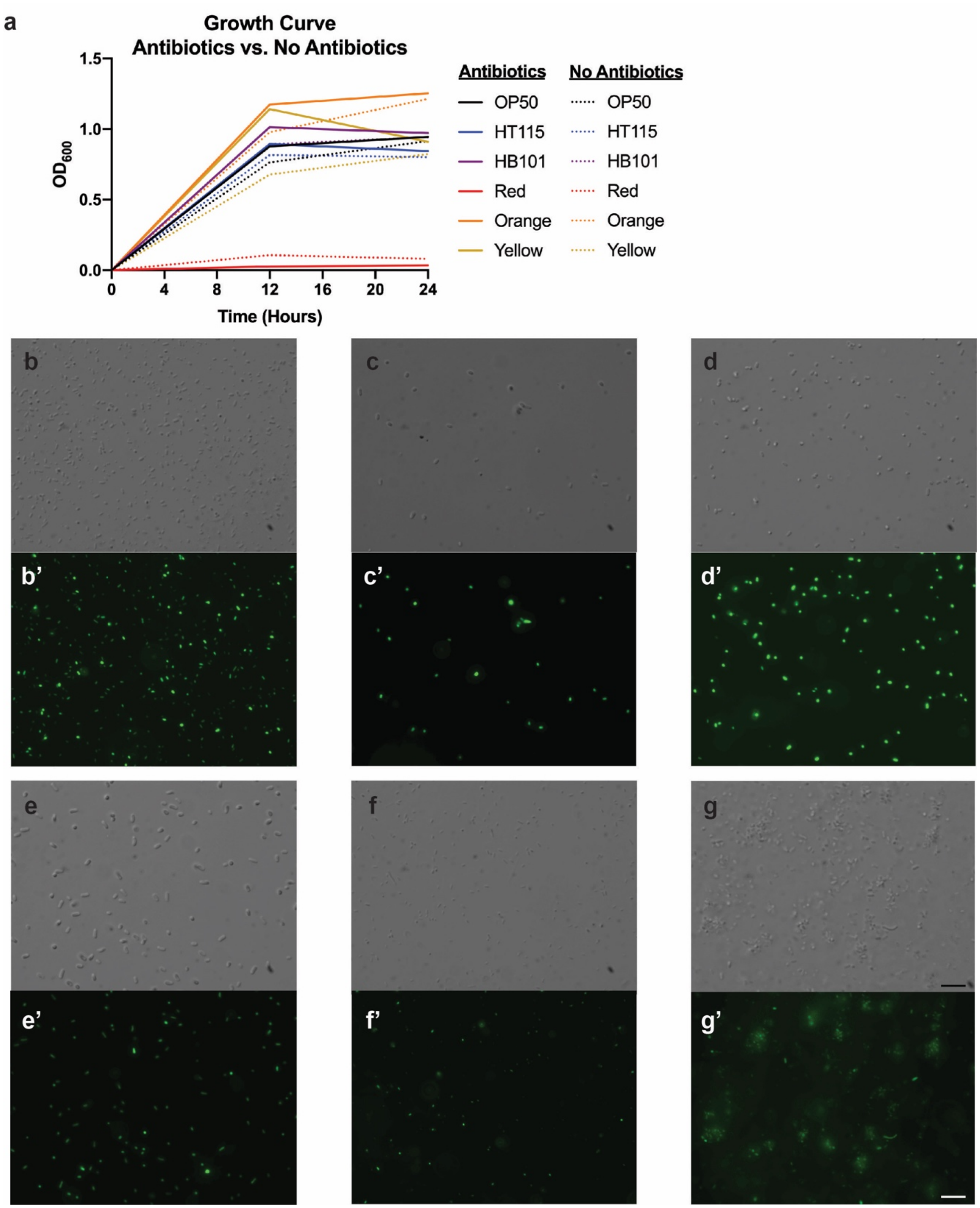


**Supplemental Figure 2. Bacterial growth curve and images.** (a) Bacterial growth curve comparison between growth in LB with and without antibiotics with no significant difference in growth rate. (b-g) Microscopic images of bacteria after being scraped off of a plate seeded at the optical density of 0.8 $A_{600}$. All bacteria are observably similar or smaller in size compared to OP50 bacteria. The bacteria are (b) OP50, (c) HT115, (d) HB101, (e) Red. (f) Orange, and (g) Yellow. (b’-g’) Images of stained bacteria with BactoView$^{TM}$ Live Fluorescent dye that stains the DNA. Scale bar 5 µm.

a

| Diet | Time to 50% of the Population Molting (Hours) | | | | Total Time to Adulthood | Hours in each molt (Average number of hours worms were GFP positive) | | | | % Faster than OP50-reared worms |
|---|---|---|---|---|---|---|---|---|---|---|
| | L1 Molt | L2 Molt | L3 Molt | L4 Molt | | L1 Molt | L2 Molt | L3 Molt | L4 Molt | |
| OP50 | 11 | 22 | 30 | 41 | 50 | 6 | 4 | 6 | 7 | - |
| HT115 | 11 | 18 | 27 | 38 | 44 | 9 | 3 | 5 | 6 | 20% |
| HB101 | 12 | 22 | 30 | 40 | 47 | 6 | 5 | 6 | 6 | 14.5% |
| Red | 11 | 20 | 28 | 38 | 44 | 9 | 4 | 4 | 6 | 20% |
| Orange | 13 | 23 | 29 | 39 | 46 | 5 | 3 | 4 | 5 | 16% |
| Yellow | 13 | 22 | 30 | 40 | 46 | 5 | 3 | 4 | 4 | 16% |

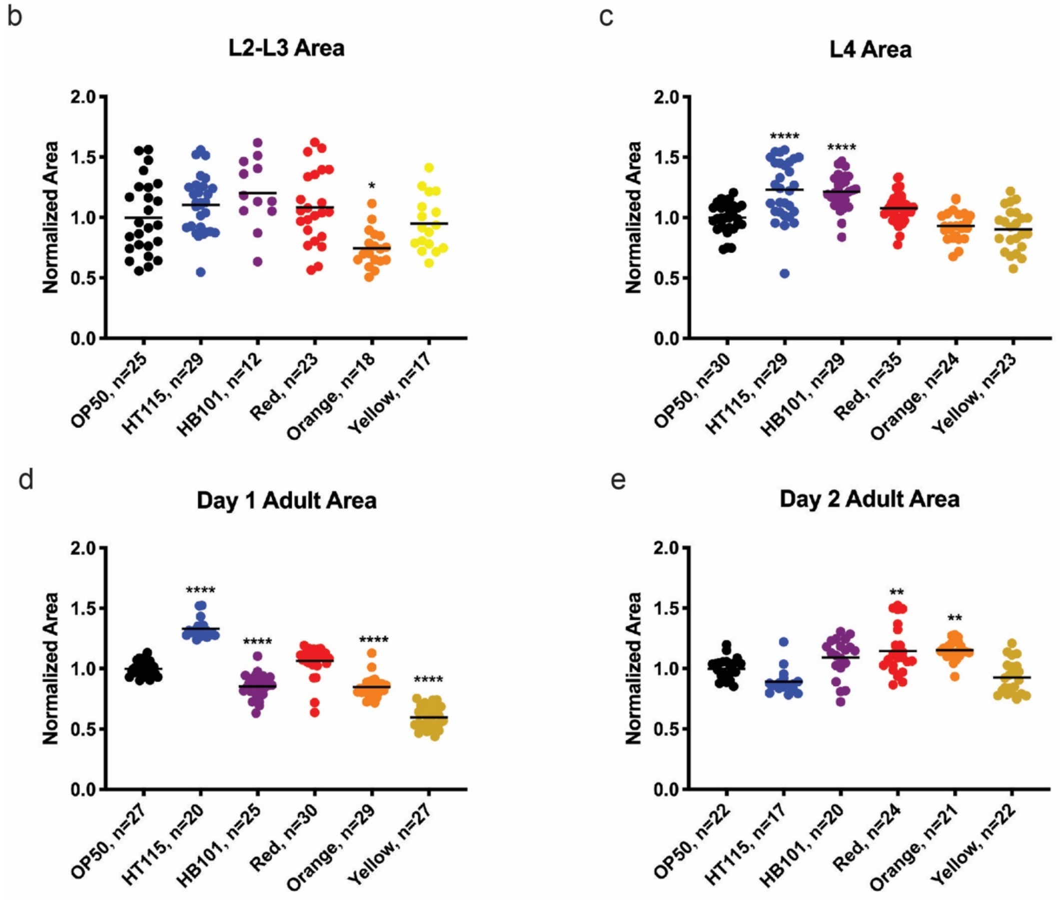


**Supplemental Figure 3. Time in each developmental stage and worm area is altered based on bacterial diet raised on.** (a) Table showing differences in time to each molt along with hours spend in each stage of development. (b-e) Area measurements of worms on each bacteria relative to OP50. (b) Orange-reared worms showed a slightly smaller area than OP50 and other bacteria. (c) *C. elegans* on HT115 and HB101 had significantly larger areas at the L4 stage. (d) Worms raised on HT115 have a larger area than worms raised on OP50 at day 1 of adulthood. HB101, Orange and Yellow worms have smaller area at day 1 of adulthood. (e) *C. elegans* raised on Red and Orange have larger areas at day 2 of adulthood compared to OP50 and the other diets. Statistical comparisons by Tukey's multiple comparison test. *, $p<0.05$; **, $p<0.01$; ***, $p<0.001$; ****, $p<0.0001$. All studies were performed in biological triplicate**.**

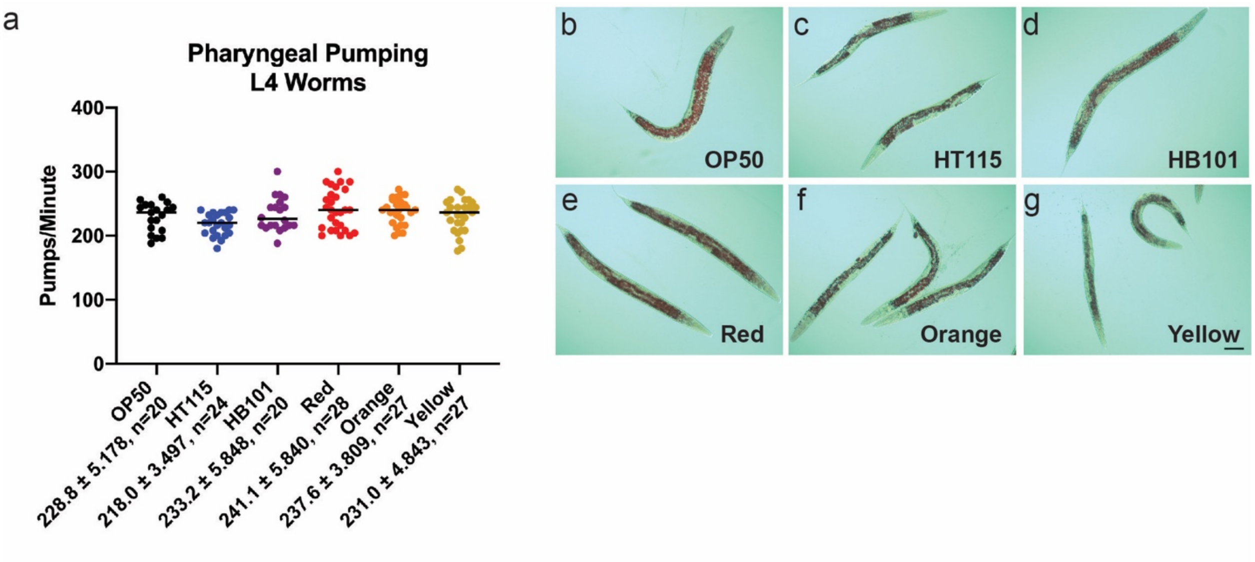


h

| | # of Colonies | |
|---|---|---|
| | 24hour | 48hour |
| OP50 | 0 | 0 |
| HT115 | 0 | 0 |
| HB101 | 0 | 0 |
| Red | 0 | 0 |
| Orange | 0 | 0 |
| Yellow | 0 | 0 |

**Supplemental Figure 4. L4 Pumping and Oil Red O staining.** (a) L4 pumping of worms raised on each bacterial diet. There was no significant difference between worms raised on the different diets. Statistical comparisons by Tukey's multiple comparison test. Study was performed in biological triplicate. (b-g) Oil Red O staining for lipid distribution in L4 *C. elegans*. Scale bar 50 μm. (h) Bacterial load experiment that counted the number of colonies that grew on an LB plate after 24 and 48 hours. n=48 individual worms per bacterial food.

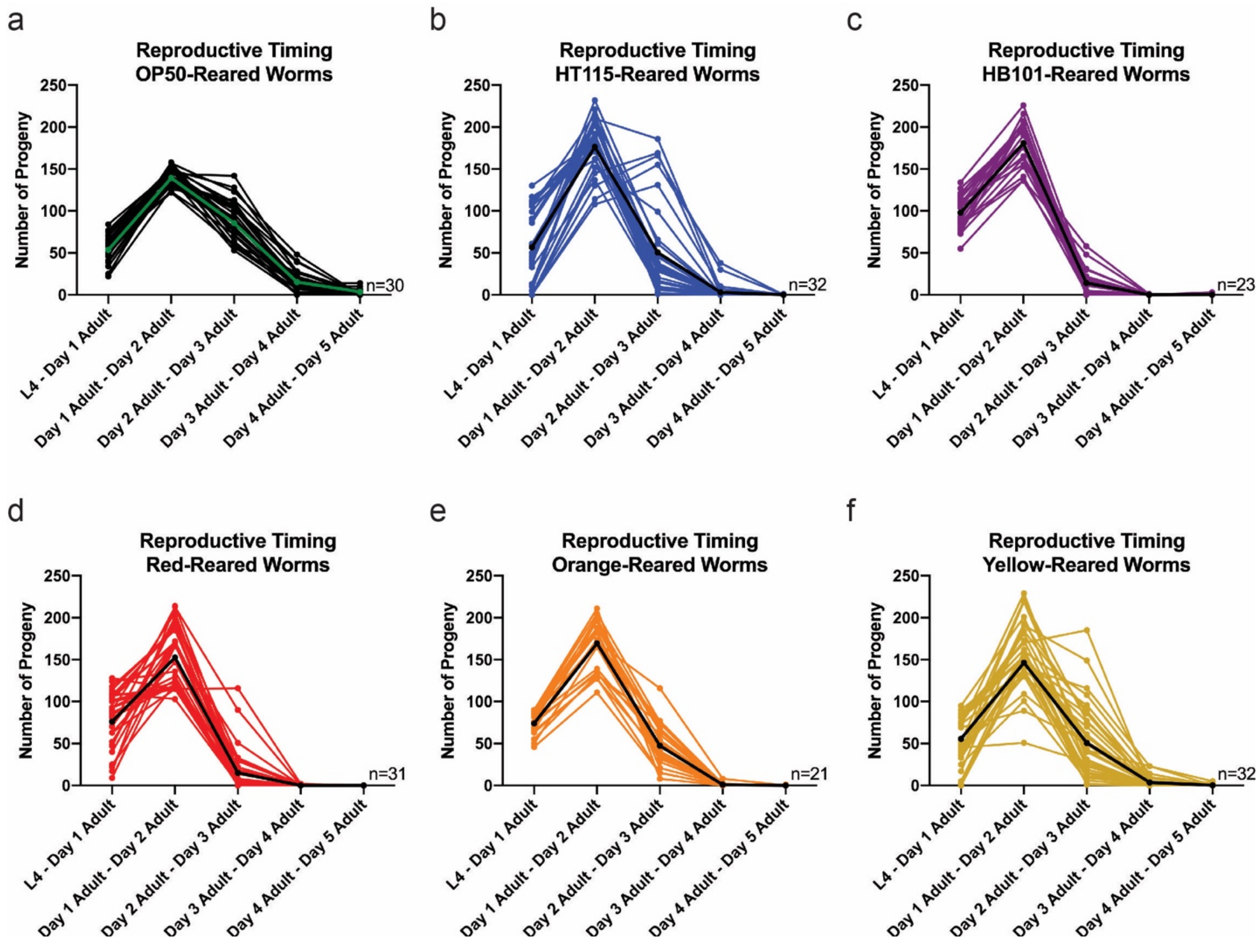


**Supplemental Figure 5 Reproductive timing is altered in *C. elegans* raised on the different bacterial diets.** Reproductive output of individual worms (each line represents a different worm) at each day of its reproductive span. The thicker green line (a) and black line (b-f) on the graph represents the average output per day of reproduction. Worms were moved every 24 hours and progeny were counted. Each bacterial food had a reproductive peak from day 1 to day 2 of adulthood and then started producing fewer progeny per day until day 5 of adulthood. HB101 (c) and Red (d) worms have a significant decline in reproductive output by day 3 of adulthood.

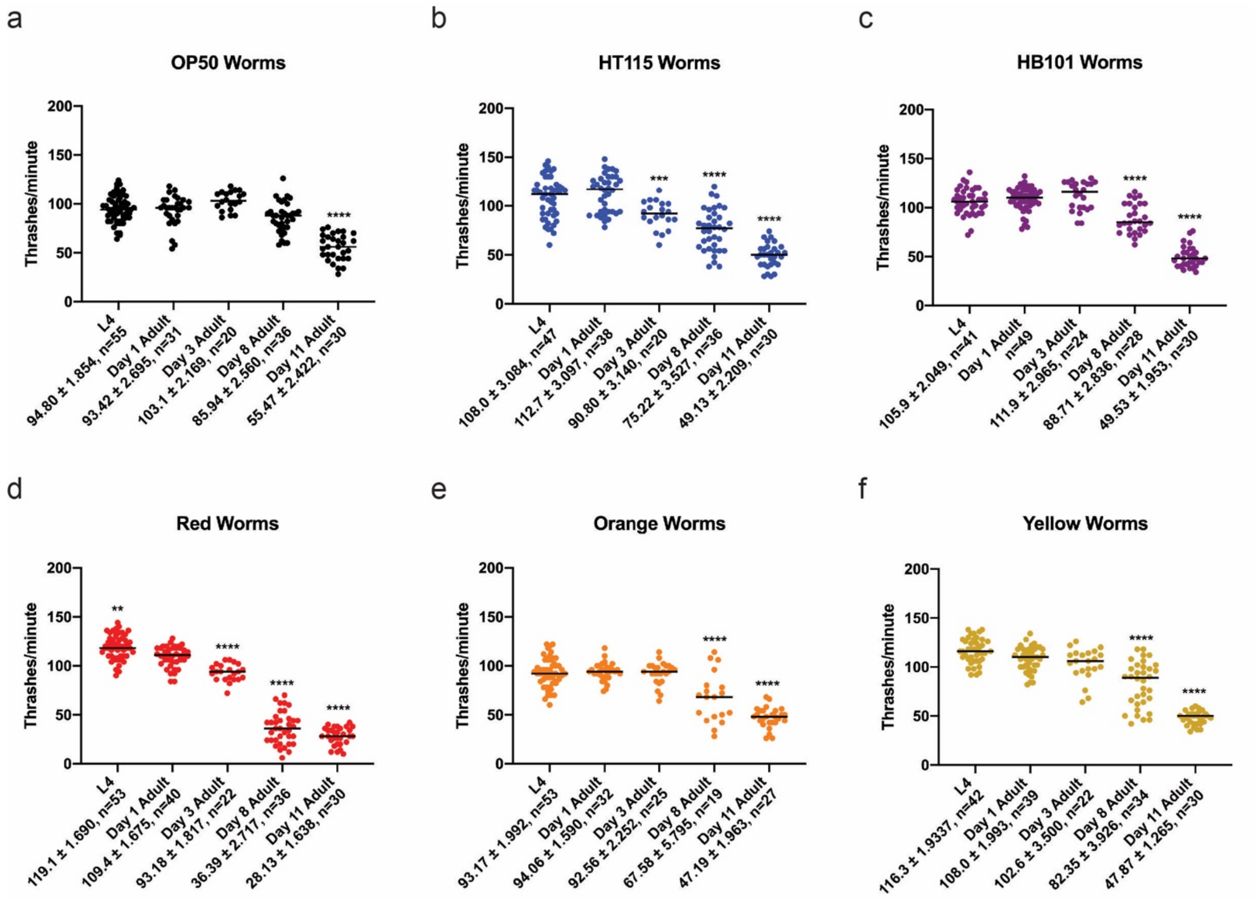


**Supplemental Figure 6 Thrashing declines with age in a food-dependent manner.** *C. elegans* at each stage of development on each bacterial diet. Comparisons were made to day 1 of adulthood. Statistical comparisons by Tukey's multiple comparison test. *, $p<0.05$; **, $p<0.01$; ***, $p<0.001$; ****, $p<0.0001$. All studies were performed in biological triplicate. (a) OP50 thrashing starts to show a decline at day 11 of adulthood. (b) HT115 thrashing declines starting at day 3 of adulthood and continues to do so up until day 11 of adulthood. (c, e-f) HB101, Orange, and Yellow decline in thrashing rate at day 8 of adulthood and continues into day 11 of adulthood. (d) Red worms decline in thrashing from day 1 to day 11 of adulthood. Red-reared worms have the most significant rate of decline in thrashing compared to the other diets.

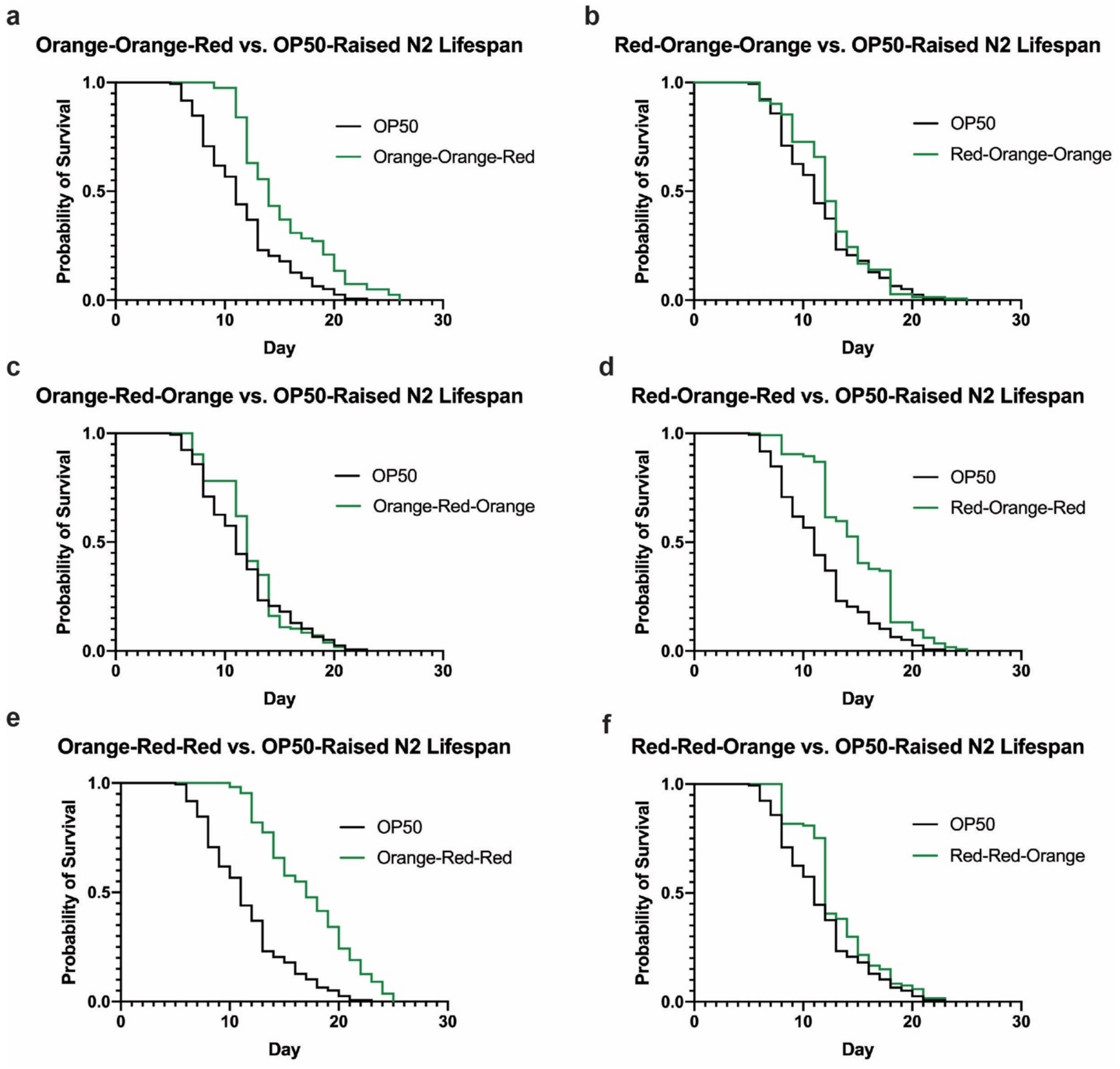


**Supplemental Figure 7 Bacterial diet combinations compared to *C. elegans* raised on OP50.** Lifespan comparisons of OP50 versus each nutraceutical diet combination. Lifespan comparisons made with Log-rank test (supplementary data 4).

### SUPPLEMENTARY TABLES

**Table S1.** Gene Ontology (GO) Terms for RNAseq analysis in L4 *C. elegans* on bacterial diets.

#### HT115-specific

**GO-TERMS UP**
organic acid metabolic process
supramolecular polymer
nucleoside phosphate binding
ribonucleotide binding
purine nucleotide binding
nucleoside phosphate metabolic process
purine nucleotide metabolic process
ribose phosphate metabolic process
actin filament-based process
lipid catabolic process
calcium ion binding
hydrolase activity acting on acid anhydrides
phosphorus metabolic process
identical protein binding
kinase binding

**GO-TERMS DOWN**
lipid catabolic process
organic acid metabolic process

#### HB101-specific

**GO-TERMS UP**
transmembrane transport
establishment of localization
metalloendopeptidase activity
purine nucleotide binding
supramolecular polymer
nucleoside phosphate binding
ribonucleotide binding
structural constituent of cuticle
peptidyl-tyrosine modification
peptidyl-serine modification

#### Red-specific

**GO-TERMS UP**
immune system process
defense response
response to biotic stimulus
phosphorus metabolic process

#### Orange-specific

**GO-TERMS DOWN**
organic acid metabolic process
lipid catabolic process

#### Yellow-specific

**GO-TERMS UP**
peptidyl-tyrosine modification
extracellular space
phosphorus metabolic process
neuropeptide signaling pathway
localization of cell
protein modification process
potassium ion transmembrane transport
peptidyl-serine modification
zinc ion binding
intrinsic component of membrane
dephosphorylation
positive regulation of kinase activity
metalloendopeptidase activity
synaptic signaling
peptidase activity

**GO-TERMS DOWN**
meiotic cell cycle
organelle fission
reproduction
chromosome segregation
cellular aromatic compound metabolic process
heterocycle metabolic process
membrane-enclosed lumen
embryo development ending in birth or egg hatching
organic cyclic compound metabolic process
ribonucleoprotein granule
RNA splicing via transesterification reactions
modification-dependent macromolecule
protein ubiquitination
organelle
covalent chromatin modification
post-embryonic animal organ development
cellular macromolecule localization
negative regulation of metabolic process
protein modification process
protein catabolic process
kinase binding
regulation of protein metabolic process
reproductive system development
post-embryonic development
development of primary sexual characteristics
identical protein binding
cellular developmental process
transcription factor complex
hydrolase activity acting on acid anhydrides
macromolecule biosynthetic process
regulation of nucleobase-containing compound
methylation
aging
small GTPase binding
process utilizing autophagic mechanism
amide transport
double-stranded DNA binding
male anatomical structure morphogenesis
oviposition
